# Simulating the Impact of Glenohumeral Capsulorrhaphy on Movement Kinematics and Muscle Function in Activities of Daily Living

**DOI:** 10.1101/2020.06.02.130880

**Authors:** Aaron S. Fox, Stephen D. Gill, Jason Bonacci, Richard S. Page

## Abstract

This study aimed to use a predictive simulation framework to examine shoulder kinematics, muscular effort and task performance during functional upper limb movements under simulated selective glenohumeral capsulorrhaphy. A musculoskeletal model of the torso and upper limb was adapted to include passive restraints that simulated the changes in shoulder range of motion stemming from selective glenohumeral capsulorrhaphy procedures (anteroinferior, anterosuperior, posteroinferior, posterosuperior, and total anterior, inferior, posterior and superior). Predictive muscle-driven simulations of three functional movements (upward reach, forward reach and head touch) were generated with each model. Shoulder kinematics (elevation, elevation plane and axial rotation), muscle cost (i.e. muscular effort) and task performance time were compared to a baseline model to assess the impact of the capsulorrhaphy procedures. Minimal differences in shoulder kinematics and task performance times were observed, suggesting that task performance could be maintained across the capsulorrhaphy conditions. Increased muscle cost was observed under the selective capsulorrhaphy conditions, however this was dependent on the task and capsulorrhaphy condition. Larger increases in muscle cost were observed under the capsulorrhaphy conditions that incurred the greatest reductions in shoulder range of motion (i.e. total inferior, total anterior, anteroinferior and total posterior conditions) and during tasks that required shoulder kinematics closer to end range of motion (i.e. upward reach and head touch). The elevated muscle loading observed could present a risk to joint capsule repair. Appropriate rehabilitation following glenohumeral capsulorrhaphy is required to account for the elevated demands placed on muscles, particularly when significant range of motion loss presents.

---

**S**urgical capsulorrhaphy is a procedure used to treat glenohumeral instability caused by elongated, lax or damaged capsule ligamentous structures.^1–6^ Glenohumeral capsulorrhaphy can be achieved by various techniques involving selective plication to different sections of the joint capsule.^1,2,4–7^ While having the primary function of correcting joint instability, the added passive tension and restricted range of motion at the shoulder may have secondary effects on active movement and muscle function. Procedures reattaching and ‘tightening’ the joint capsule will invariably impact glenohumeral joint motion and forces by altering the passive resistance of the ligaments around the joint.^1,8,9^ Variable restrictions to range of motion have been observed under differing patterns of localised plication.^1^ The restricted range of motion following surgical glenohumeral capsulorrhaphy may make achieving shoulder postures required for activities of daily living difficult for patients. Further, muscular effort may need to increase during such tasks in order to counter the added passive resistance from a ‘tighter’ joint capsule. Understanding the impact of the technical variations in glenohumeral capsulorrhaphy on movement and muscle function is therefore relevant to the design of rehabilitation strategies and return to function following the procedure. Knowledge of how movement may be altered, and how muscle function is affected will assist in providing specific rehabilitation targets.

Various studies^10–14^ have used musculoskeletal simulations built from experimental movement data to examine how changes in the musculoskeletal system impact muscle and/or joint loading. Within these studies – parameters in the musculoskeletal model are altered to represent relevant clinical pathologies, with the altered model/s then used in simulations which estimate the neuromuscular control strategies (i.e. muscle activation and forces) that reproduce the experimental data. This approach has been applied in a gait analysis context,^10–12^ to examine the effects of lower limb^10^ and deep core^11^ muscle weakness on neuromuscular control and joint loading during walking and running gait, respectively. Similarly, the effect of simulated hip strengthening exercises on hip joint forces during walking has been explored in patients with total hip arthroplasty.^12^ Musculoskeletal simulations of individuals with cerebral palsy have also been conducted to understand how the common neuromuscular deficits associated with the condition (i.e. muscle weakness and contracture) impact gait function.^13,14^ Through appropriate musculoskeletal simulation, these studies have identified how deficits or changes in the musculoskeletal system may impact muscle function and/or joint loading – providing relevant clinical targets for rehabilitation strategies.

A limitation of these studies is their use of experimental kinematic data to generate muscle-driven simulations. During these simulations, the movement strategy is held consistent in order to match the experimental data. This approach can reveal information about how neuromuscular strategies may adapt with specific changes in the musculoskeletal system – but is limited in its ability to identify how movement may change, and how movement changes would affect the neuromuscular strategies used. This may not reflect what naturally occurs with musculoskeletal system changes. For example, substantial differences in kinematic strategies during activities of daily living have been observed between elderly individuals with symptomatic rotator cuff impingements versus asymptomatic individuals;^15^ as well as patients with glenohumeral osteoarthritis and those who have undergone total shoulder arthroplasty compared to healthy controls.^16^ Predictive simulations that are generated *de novo* (i.e. without experimental data) can assist in identifying cause and effect relationships between changes to musculoskeletal parameters, and the movement and neuromuscular strategies used. Predictive simulations typically provide ‘goals’ and/or constraints on a task, and the optimal movement and neuromuscular strategies that meet these are determined. Changes in musculoskeletal parameters may alter the optimal movement and/or neuromuscular strategy, and this is therefore captured through the predictive simulation. Similar to musculoskeletal simulations, the predictive simulation approach has been more readily applied in a gait context.^17,18^ Song and Geyer^18^ induced neural, muscular and skeletal deficits associated with ageing to generate predictive simulations of walking in the elderly. Muscle strength and mass were subsequently identified as the predominant factors responsible for the typical gait changes seen in elderly populations.^18^ Ong et al.^17^ simulated isolated weakness or contracture of the plantarflexor muscles to emulate adaptations in the musculoskeletal system typically observed with cerebral palsy. Muscle weakness elucidated a slower ‘heel-walking’ gait, while contracture resulted in a crouched ‘toe-walking’ gait – agreeing with the common gait adaptations seen in experimental evaluations of those with cerebral palsy.^17^ These studies reveal the power of predictive simulation for investigating how movement and control strategies change with adaptations in the musculoskeletal system.

Musculoskeletal modelling and predictive simulation can be used to investigate the effects of selective glenohumeral capsulorrhaphy associated with capsulolabral repair. This framework provides an opportunity to alter parameters in a musculoskeletal model that simulate the changes in passive glenohumeral joint resistance observed with selective glenohumeral capsulorrhaphy – and examine how these changes impact movement and muscle function during upper limb tasks. These findings can prove useful in providing surgeons with an understanding of the impact their procedures may have on daily function, and subsequently promote a more targeted approach to rehabilitation following selective glenohumeral capsulorrhaphy – whereby exercises that incorporate the most affected muscles could be targeted. This study aimed to use a predictive simulation framework to examine shoulder kinematics, muscular effort and task performance time during functional upper limb movement tasks under various simulated selective glenohumeral capsulorrhaphy conditions. We systematically manipulated passive restraints at the glenohumeral joint within a musculoskeletal model to simulate the impact of different localised glenohumeral joint plications. It was hypothesised that the selective capsulorrhaphy conditions would: (i) alter the movement strategies used; (ii) increase the muscular effort; and (iii) reduce performance (i.e. increased time to complete the task).

## Methods

### Musculoskeletal Model

A generic seven-segment (torso, clavicle, scapula, humerus, ulna, radius and hand) musculoskeletal model of the torso and right upper limb was developed in OpenSim (version 4.0)^19^ and used in this study. The segment properties and inertial parameters used reflected those in the model provided by Wu et al.^20^. The kinematic foundation for the model^21,22^ included five degrees of freedom (DOF) describing the kinematics of the shoulder girdle, elbow and forearm as recommended by the International Society of Biomechanics.^23^ The shoulder included three DOFs describing the elevation plane, elevation angle and shoulder axial rotation.^21,22^ Certain kinematic ranges were expanded from the original kinematic model^21,22^ to permit the motions observed in experimental studies of the functional tasks being simulated. Specifically, the elevation plane range was expanded to −95°to 130°and humeral axial rotation was expanded to −90°(external rotation) to 130°(internal rotation).^24^ Overall motion of the shoulder girdle (including the clavicle and scapula) was determined by the regression equations described by de Groot and Brand^25^, and driven by the shoulder elevation angle. The elbow and forearm included one DOF each describing elbow flexion and pronation/supination, respectively.^21,22^ Coordinate limit forces were linked to each shoulder DOF in the model that applied a constant damping force equivalent to 0.1 Nms·rad^-1^ throughout the range of motion.^22^ Motion of the torso was locked and lower limb segments were not included to ensure the simulated tasks were achieved by upper limb motion alone. The wrist and finger joints were also locked to minimise the DOFs of the model, reducing simulation complexity and computation times. The model was actuated by 26 Hill-type muscle-tendon units representing the major axio-scapular, axio-humeral and scapulo-humeral muscles.^20^

The muscle model parameters were set to those used in De Groote et al.^26^ and tendon dynamics were ignored. The muscle-tendon paths and wrapping points of the model have been optimised to match muscle moment arms measured in cadaveric upper extremities;^20^ while the muscle-tendon unit parameters were derived from an optimisation routine from sets of isometric and isokinetic tasks performed on one generic healthy individual.^20^ Idealised torque actuators were used to drive elbow and forearm motion.

### Models of Selective Capsulorrhaphy

Our study created models simulating selective glenohumeral capsulorrhaphy. A baseline model of no joint capsulorrhaphy was created (i.e. ‘None’ model), followed by models representing anteroinferior, anterosuperior, posteroinferior, posterosuperior, total anterior, total inferior, total posterior and total superior glenohumeral capsulorrhaphy procedures.^1^ The models were based off data presented by Gerber et al.^1^ – whereby the passive restraints included in the model were optimised so that similar passive joint angles were reached with identical applied torques as observed in their experiments. The shortening of the shoulder capsule achieved through capsulorrhaphy typically reduces glenohumeral joint range of motion.^1^ To appropriately represent this in our musculoskeletal model, force generating objects representing the passive restraints (i.e. ligaments, joint capsule) of the glenohumeral joint that responded to joint angle needed to be included. Gerber et al.^1^ demonstrated that passive glenohumeral range of motion is impacted by combined joint positions (e.g. shoulder axial rotation range varies as shoulder elevation increases). We adapted the original *Expression Based Coordinate Force* into a custom plugin (named as the *Dual Expression Based Coordinate Force*) to account for this behaviour. This custom force class uses an expression (i.e. mathematical formula) with two joint angles (e.g. shoulder axial rotation and elevation) as inputs to calculate the restrictive force to be applied to the relevant joint motion. Two *Dual Expression Based Coordinate Force* objects were included in each model – one that restricted shoulder elevation relative to shoulder elevation angle and elevation plane; and one that restricted shoulder axial rotation relative to shoulder axial rotation and shoulder elevation.

An optimisation routine was used to identify the mathematical expressions required to accurately model the passive restraints under different capsulorrhaphy conditions. A simplified version of the musculoskeletal model containing only the humerus and scapula with all muscles disabled, and *Coordinate Limit Forces* placed on the shoulder axial rotation and elevation coordinates was developed. The choice to use the more basic *Coordinate Limit Force* class at this stage was to simplify the optimisations, as only one joint angle was altered in each simulation. The model was placed in identical starting positions to those described by Gerber et al.^1^, and identical torques (i.e. 0.5 and 1.0 Nm for axial rotation and elevation motions, respectively) were then applied to the humerus. The lower limit (i.e. the joint angle where resistive force begins to develop) and stiffness (i.e. the rotational stiffness of the passive force when the joint angle exceeds the lower limit) parameters of the *Coordinate Limit Forces* were optimised so that an appropriate magnitude of resistive force was applied to limit the joint motion to the peaks recorded by Gerber et al.^1^. This process was repeated to identify optimised resistive force application across the range of joint motions at specific joint positions (i.e. abduction; flexion; and internal and external rotation at zero, 45 and 90 degrees of abduction) for each of the capsulorrhaphy conditions (i.e. none; anteroinferior; anterosuperior; posteroinferior; posterosuperior; total anterior; total inferior; total posterior; total superior). To identify the mathematical expression (i.e. the input to the *Dual Expression Based Coordinate Force*) that fit the optimised passive resistive forces, the resistive force generated by the *Coordinate Limit Forces* in each simulation were plotted against joint motion. A one-term Gaussian exponential model, as specified by:

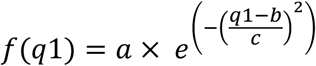

where *q*1 referred to the joint angle being tested (e.g. shoulder elevation or shoulder axial rotation), and *a, b*, and *c* referred to equation coefficients was then fit to the resistive force-joint motion data from each simulation. The one-term Gaussian exponential model was selected as the most appropriate fit option as it produced high R^2^ values (i.e. greater than 0.99) and minimised the root mean square error between the curve fit and resistive force-joint motion data in comparison to other available mathematical models. The process resulted in a Gaussian exponential model being fit for each joint position tested. To finalise the expression to provide in the model’s two *Dual Expression Based Coordinate Force* objects, the change in the coefficient values (i.e. *a, b*, and *c*) with different joint positions for the joint angle being tested were modelled against specific equations. For the force responsible for restricting shoulder elevation relative to shoulder elevation angle and elevation plane – two joint positions (i.e. the abduction and flexion elevation planes) were tested. Having only two positions meant a linear fit was the only option for modelling the coefficients in this expression. Subsequently, the coefficients (i.e. *a, b*, and *c*) for this force were specified by:

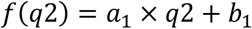

where *q*2 referred to the joint position being tested (i.e. elevation plane), and *a*_1_ and *b*_1_ referred to equation constants that were optimised to ensure the curve fit the data.

For the force responsible for restricting shoulder axial rotation relative to shoulder axial rotation and shoulder elevation – three joint positions (i.e. 0°, 45°and 90°of shoulder elevation) were tested. A quadratic equation was found to provide the best fit for these data. Subsequently, the coefficients (i.e. *a, b*, and *c*) for this force were specified by:

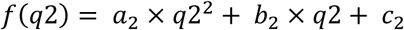

where *q*2 referred to the joint position being tested (i.e. shoulder elevation), and *a*_2_, *b*_2_, and *c*_2_ referred to equation constants that were optimised to ensure the curve fit the data.

### Predictive Simulations

Each capsulorrhaphy model was used in predictive muscle-driven simulations of functional movements (see Table 1 and Online Videos <u>1</u>, <u>2</u> & <u>3</u>) generated using OpenSim Moco (version 0.3.0)^27^ in MATLAB (version 2019b, The Mathworks Inc., Natick, MA, United States). OpenSim Moco provides a framework for solving optimal control problems for musculoskeletal systems using direct collocation. The predictive simulations generate the control signals (i.e. muscle activations) required to produce the desired movement.

**Table 1.**
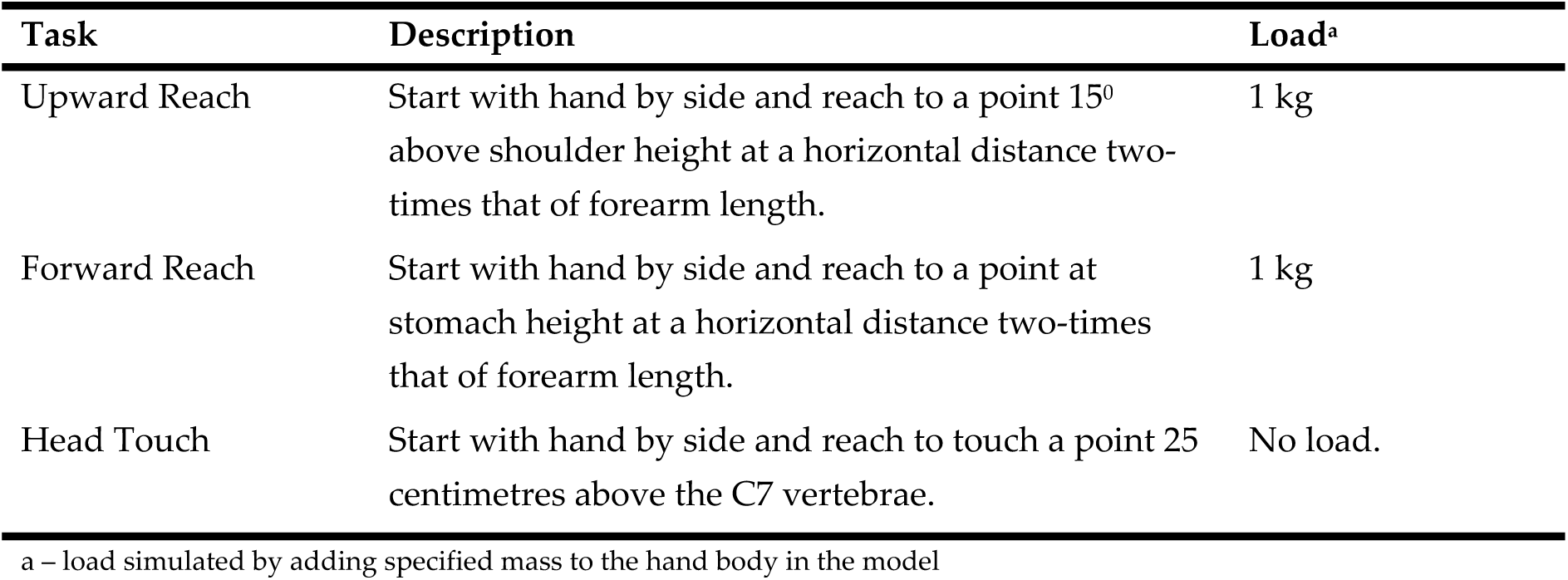
Description of functional movements simulated using OpenSim Moco

The functional movements were designed to replicate experimental tasks measured in the literature.^24,28^ The predictive simulations were generated by prescribing a series of task constraints and goals that contributed to an objective function value, with the optimal control problem aiming to minimise this value. First, all tasks were constrained to begin with the model set in a stationary neutral position with the arm placed by the side (i.e. 0°of shoulder elevation and rotation, elbow flexion, and pronation/supination). Second, upper and lower bounds were placed on the shoulder joint angles by extracting the maximum and minimum values recorded in similar movement tasks^24,28,29^ – to ensure that the desired movements were achieved with relatively ‘normal’ kinematic strategies. Third, the joint angular velocities were constrained to be zero at the end of the movement to ensure the limb decelerated and finished in a static position. Fourth, a point (or set of points) on the model were provided with an end-point goal relevant to the desired movement (i.e. the optimal solution aimed to minimise the distance of the point(s) on the model to the desired end-point to ensure the hand reached the desired end position relevant to the task). Fifth, a goal to minimise the sum of squared control signals (i.e. muscle activations and torque actuators) was included as an effort criterion (i.e. minimising effort). Sixth, a goal to minimise the time taken to reach the desired movement end-point was included as a performance criterion. Overall, this generated predictive simulations of the tasks that achieved the desired movement (i.e. reaching the end task goals while remaining within the kinematic bounds), while minimising effort (i.e. neuromuscular strategy used) and maximising performance (i.e. time taken to perform the task). Predictive simulations of each movement task were repeated across each of the capsulorrhaphy models, resulting in a total of 27 predictive simulations (i.e. 3 tasks x 9 models).

An important step in solving optimal control problems is determining an appropriate mesh interval (i.e. time step) and, subsequently, the number of nodes (i.e. ‘node’ or ‘grid density,’ or sample points) to use for the solution. We used a ‘grid refinement’ approach as outlined in Lee and Umberger^30^ to determine an appropriate node density for each of the movement tasks. Specifically, this approach solves optimal control problems for the task with an increasing number of nodes. An appropriate node density is identified where the value of the objective function begins to plateau, and minimal changes in the model outputs are observed as a result of increasing node number.^30^ The duration of time taken to solve the optimal control problem must also be considered in the context of the added benefit a denser grid may provide.^30^ We examined the outputs of solutions at different node densities, and plotted the root mean square error of the model outputs at adjacent node densities to identify where similar results began to present. Node densities of 25, 51, 101, 151, 201 and 251 were tested for each of the movement tasks using the ‘None’ model. The solution of the previous node density was used as the initial guess for the subsequent problem to speed up computation time.^30^ Similarly, computation time for the optimal control problems using the remaining capsulorrhaphy models was reduced by using the relevant node density solution for the ‘None’ model as the initial guess.

### Data Analysis

The effect of capsulorrhaphy on shoulder function was assessed by comparing the shoulder kinematics and a metric of muscle cost, along with the general performance indicator of time taken to complete the task between the predictive simulations generated by the different capsulorrhaphy models. Only a single model was simulated for each capsulorrhaphy condition, therefore descriptive rather than inferential comparisons were made. Shoulder elevation, axial rotation and elevation plane angles were directly extracted from the optimal control solutions. Joint angles were time-normalised (i.e. 0%-100% reflecting the start- and end-points of the movement, respectively), and the results from each capsulorrhaphy model in each task were visually compared to the ‘None’ model as a reference. Mean absolute errors across the shoulder joint angles were calculated for the selective capsulorrhaphy conditions relative to the ‘None’ model to examine the magnitude of difference in joint kinematics during the simulated tasks.

Individual and total muscle costs^10^ were calculated and descriptively compared for each of the predictive simulations. An increase in muscle cost denotes an increased load on the muscle(s) which (in the absence of adequate recovery) may lead to fatigue, damage or muscle weakening.^10^ Individual muscle costs (*M*_*cost*_) (i.e. the ratio of force produced by the muscle relative to its potential maximum) was quantified for each muscle at each simulation time-step as per van der Krogt et al.^10^, using:

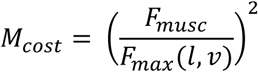

where *F*_*musc*_ is the muscles current level of force output, and *F*_*max*_(*l, v*) is the instantaneous maximum muscle force achievable considering the current length and velocity of the muscles fibres. Total muscle cost was calculated as the sum of all individual muscle costs integrated over time.^10^ The percentage change in total muscle cost was calculated for the selective capsulorrhaphy conditions relative to the ‘None’ model. The relative contribution of individual muscles to the relative change in total muscle cost across the simulated capsulorrhaphy conditions were calculated to determine the muscles primarily responsible for the observed changes.

All code used to generate the models and simulations, along with the resulting data are available at https://simtk.org/projects/gh-caps-sims.

## Results

### Capsulorrhaphy Models

Under the simulated passive range of motion test conditions, all models fell within 0.1 degrees of the data presented by Gerber et al.^1^ (see Table 2). Reductions in range of motion were achieved via resistive force being applied earlier during movement (see Figures 1 and 2).

**Table 2.** Simulated maximal range of motion values for selective capsulorrhaphy models compared to those obtained by Gerber et al.^1^ (in brackets)

| Capsulorrhaphy Model | Abduction | Flexion | Ext. Rot.<br>(0° Abd.) | Int. Rot.<br>(0° Abd.) | Ext. Rot.<br>(45° Abd.) | Int. Rot.<br>(45° Abd.) | Ext. Rot.<br>(90° Abd.) | Int. Rot.<br>(90° Abd.) |
| --- | --- | --- | --- | --- | --- | --- | --- | --- |
| None | 91.50<br>(91.5) | 85.52<br>(85.6) | 53.37<br>(53.4) | 44.59<br>(44.6) | 104.33<br>(10.4) | 38.99<br>(39.0) | 132.96<br>(133.0) | 30.74<br>(30.8) |
| Anteroinferior | 72.16<br>(72.1) | 70.54<br>(70.5) | 32.74<br>(32.8) | 44.07<br>(44.1) | 69.28<br>(69.3) | 36.03<br>(36.1) | 87.23<br>(87.3) | 23.06<br>(23.1) |
| Anterosuperior | 89.74<br>(89.8) | 71.75<br>(71.8) | 23.28<br>(23.3) | 44.07<br>(44.1) | 85.74<br>(85.8) | 36.34<br>(36.4) | 123.86<br>(123.9) | 26.88<br>(26.9) |
| Posteroinferior | 80.05<br>(80.0) | 71.44<br>(71.5) | 49.54<br>(49.6) | 35.54<br>(35.6) | 103.92<br>(104.0) | 21.39<br>(21.4) | 131.04<br>(131.1) | 10.29<br>(10.3) |
| Posterosuperior | 82.02<br>(82.1) | 76.71<br>(76.8) | 48.83<br>(48.9) | 28.43<br>(28.5) | 103.33<br>(103.4) | 28.24<br>(28.3) | 130.74<br>(130.8) | 27.49<br>(27.5) |
| Total Anterior | 73.85<br>(73.9) | 66.32<br>(66.3) | 21.27<br>(21.3) | 42.72<br>(42.8) | 67.24<br>(67.3) | 35.23<br>(35.3) | 86.06<br>(86.1) | 21.38<br>(21.4) |
| Total Inferior | 63.82<br>(63.8) | 65.82<br>(65.8) | 32.74<br>(32.8) | 37.49<br>(37.5) | 74.54<br>(74.6) | 19.18<br>(19.2) | 87.23<br>(87.3) | 3.99<br>(4.0) |
| Total Posterior | 76.64<br>(76.6) | 66.50<br>(66.5) | 49.07<br>(49.1) | 23.04<br>(23.1) | 107.75<br>(107.8) | 11.77<br>(11.8) | 129.45<br>(129.5) | 9.80<br>(9.80) |
| Total Superior | 89.95<br>(90.0) | 68.26<br>(68.3) | 17.80<br>(17.8) | 28.98<br>(29.0) | 82.28<br>(82.3) | 27.78<br>(27.8) | 115.44<br>(115.5) | 24.58<br>(24.6) |

Ext. – external; Int. – internal; Rot. – rotation; Abd. – abduction

**Figure 1.**
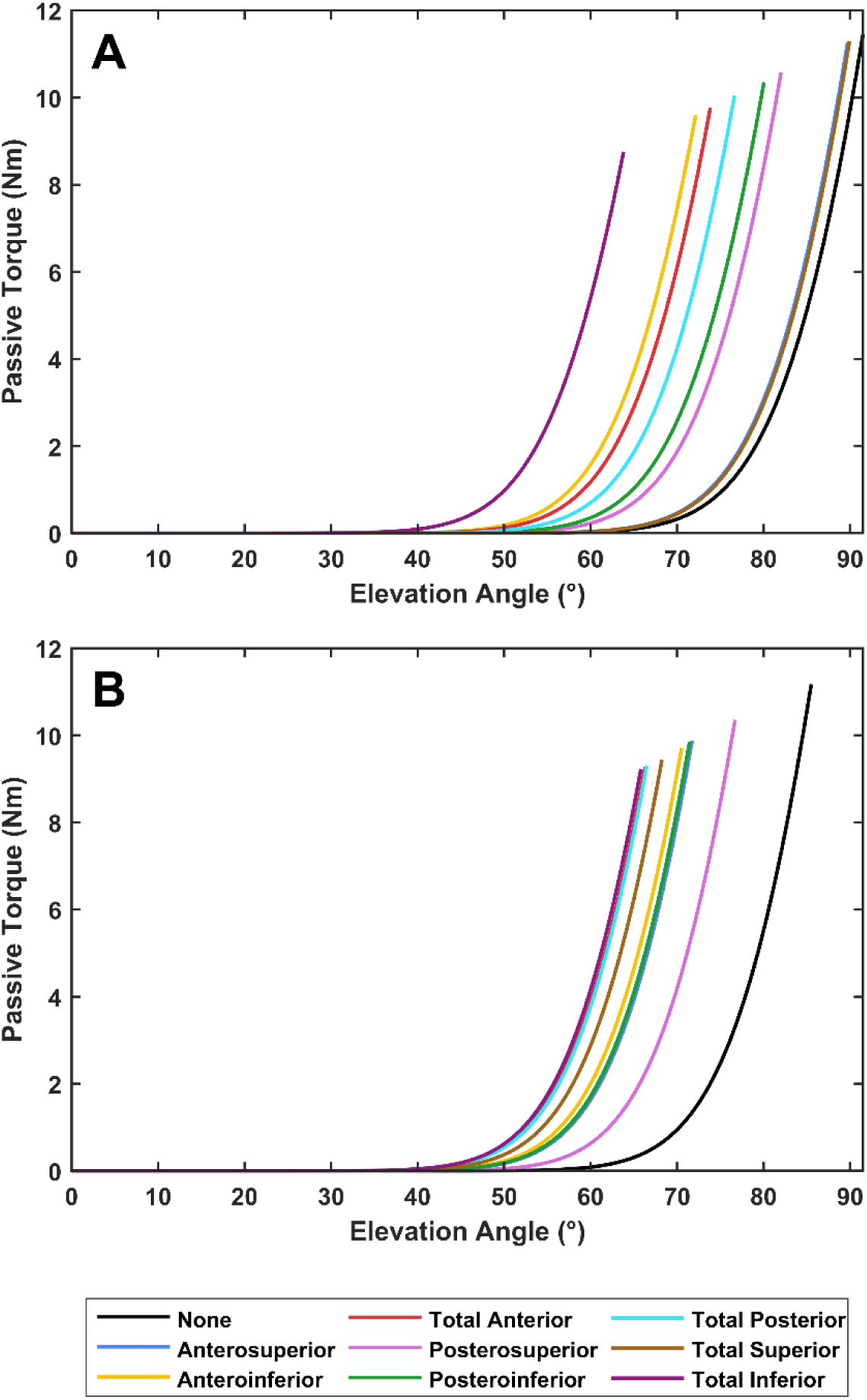
Passive torque generated by the simulated passive restraints through shoulder elevation in the abduction (A) and flexion (B) elevation planes for the baseline (i.e. None) and capsulorrhaphy models. Each curve terminates at the maximal range of motion for a model.

**Figure 2.**
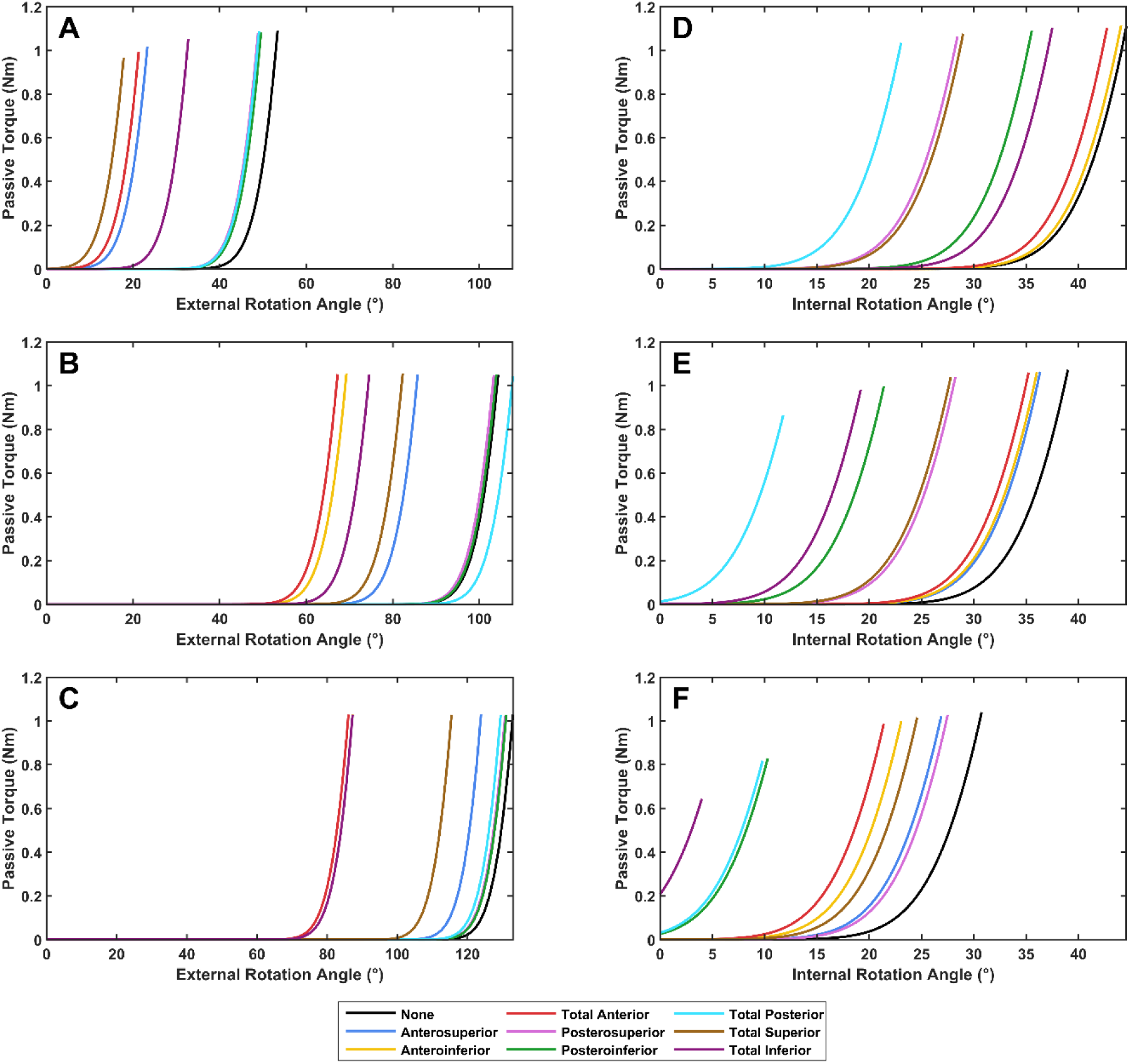
Passive torque generated by the simulated passive restraints through shoulder axial external (A, B and C) and internal rotation (D, E and F) at 0, 45 and 90 degrees of shoulder elevation in the abduction plane, respectively, for the baseline (i.e. None) and capsulorrhaphy models. Each curve terminates at the maximal range of motion for a model.

### Grid Refinement

Based on the grid refinement approach – node densities of 201, 201 and 101 were selected for the upward reach, forward reach and head touch tasks, respectively. A full summary of the grid refinement results are presented in Supplementary Document 1.

### Shoulder Kinematics

Joint angles for each of the selective capsulorrhaphy models across the three simulated tasks are presented in Figures 3-5. There were minimal differences in shoulder elevation between the selective capsulorrhaphy models compared to the ‘None’ model, with mean absolute error typically less than two degrees (see Table 3). An exception to this was in the head touch task, where the error in shoulder elevation was larger (i.e. ∼3-4 degrees) in the anteroinferior, posteroinferior, posterosuperior, total anterior and total posterior conditions (see Table 3). This appeared to be mostly driven by a reduction in shoulder elevation throughout the entirety of the task (see Figure 5). The elevation angle was highly consistent across all tasks and capsulorrhaphy conditions, with all mean absolute errors under 0.75 degrees (see Table 3). The consistency of shoulder axial rotation angles was task dependent. Shoulder axial rotation angles were highly consistent across the capsulorrhaphy conditions in the forward reach task, with all mean absolute errors under 0.42 degrees (see Table 3). Larger mean absolute errors (i.e. ∼1-2 degrees) were observed for shoulder axial rotation angles across the majority of capsulorrhaphy conditions for the head touch task, with the exception of the total posterior condition (4.50 degrees) (see Table 3) – driven by a reduction in shoulder internal rotation during the task (see Figure 5). Small mean absolute errors (i.e. < 0.7 degrees) were observed for shoulder axial rotation angles across the anteroinferior, anterosuperior and total anterior conditions for the upward reach task; while larger errors (i.e. ∼2-5 degrees) were observed across the remaining capsulorrhaphy conditions (see Table 3).

**Table 3.** Mean absolute error for shoulder joint angles (in degrees) relative to the ‘None’ model during the simulated tasks across the selective capsulorrhaphy models.

| Capsulorrhaphy Model | Upward Reach |  |  | Forward Reach |  |  | Head Touch |  |  |
| --- | --- | --- | --- | --- | --- | --- | --- | --- | --- |
|  | Shoulder Elevation | Elevation Angle | Axial Rotation | Shoulder Elevation | Elevation Angle | Axial Rotation | Shoulder Elevation | Elevation Angle | Axial Rotation |
| Anteroinferior | 1.26 | 0.16 | 0.58 | 0.18 | 0.04 | 0.24 | 3.58 | 0.02 | 1.63 |
| Anterosuperior | 1.00 | 0.07 | 0.67 | 0.08 | 0.02 | 0.18 | 0.24 | 0.01 | 0.12 |
| Posteroinferior | 1.19 | 0.75 | 2.32 | 0.27 | 0.10 | 0.39 | 3.70 | 0.23 | 1.62 |
| Posterosuperior | 1.11 | 0.34 | 3.36 | 0.26 | 0.06 | 0.42 | 3.45 | 0.30 | 1.18 |
| Total Anterior | 1.31 | 0.22 | 0.37 | 0.17 | 0.03 | 0.42 | 3.91 | 0.03 | 1.47 |
| Total Inferior | 1.50 | 0.63 | 2.73 | 0.27 | 0.11 | 0.35 | 1.69 | 0.53 | 1.65 |
| Total Posterior | 1.62 | 0.65 | 5.23 | 0.40 | 0.17 | 0.39 | 4.18 | 0.07 | 4.50 |
| Total Superior | 1.47 | 0.25 | 3.52 | 0.26 | 0.06 | 0.41 | 0.05 | 0.03 | 0.79 |

**Figure 3.**
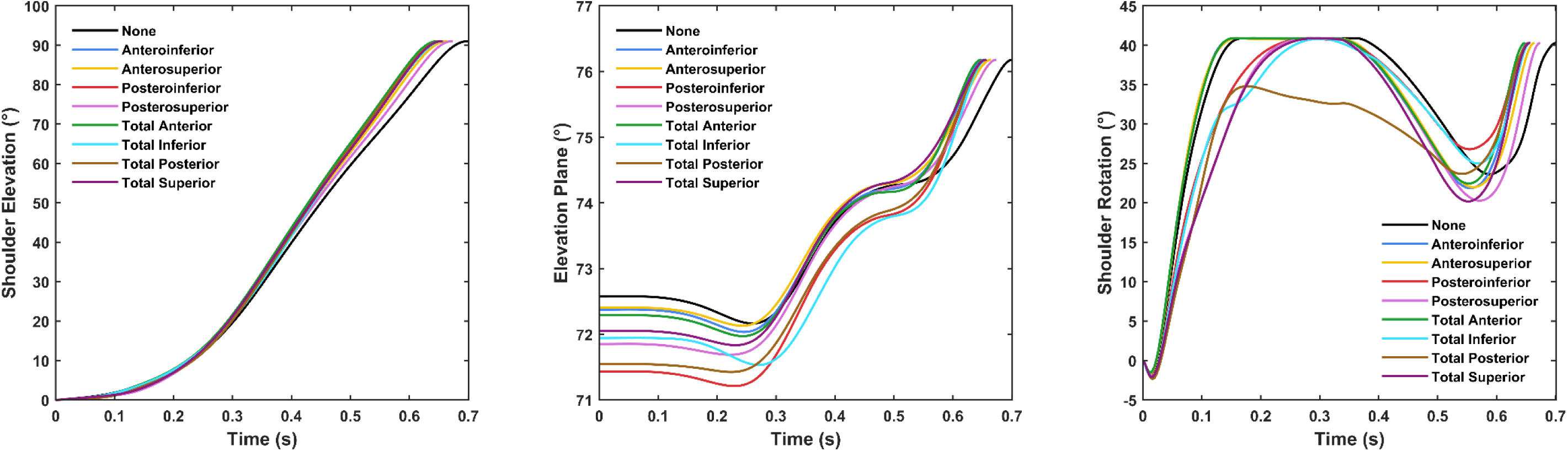
Shoulder kinematics for the baseline (i.e. None) and simulated capsulorrhaphy models during the upward reach task. More positive elevation plane values signify elevation shifting further in front of the body (i.e. 90 degree elevation plane refers to pure sagittal plane flexion). Positive and negative shoulder axial rotation values refer to internal and external rotation, respectively.

**Figure 4.**
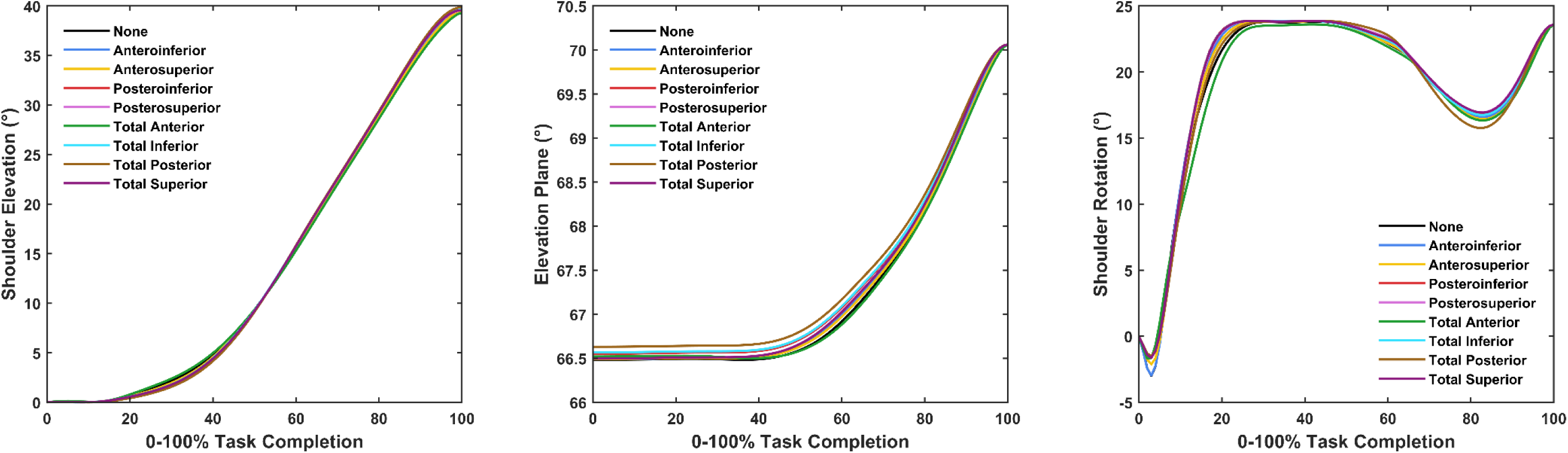
Shoulder kinematics for the baseline (i.e. None) and simulated capsulorrhaphy models during the forward reach task. More positive elevation plane values signify elevation shifting further in front of the body (i.e. 90 degree elevation plane refers to pure sagittal plane flexion). Positive and negative shoulder axial rotation values refer to internal and external rotation, respectively.

**Figure 5.**
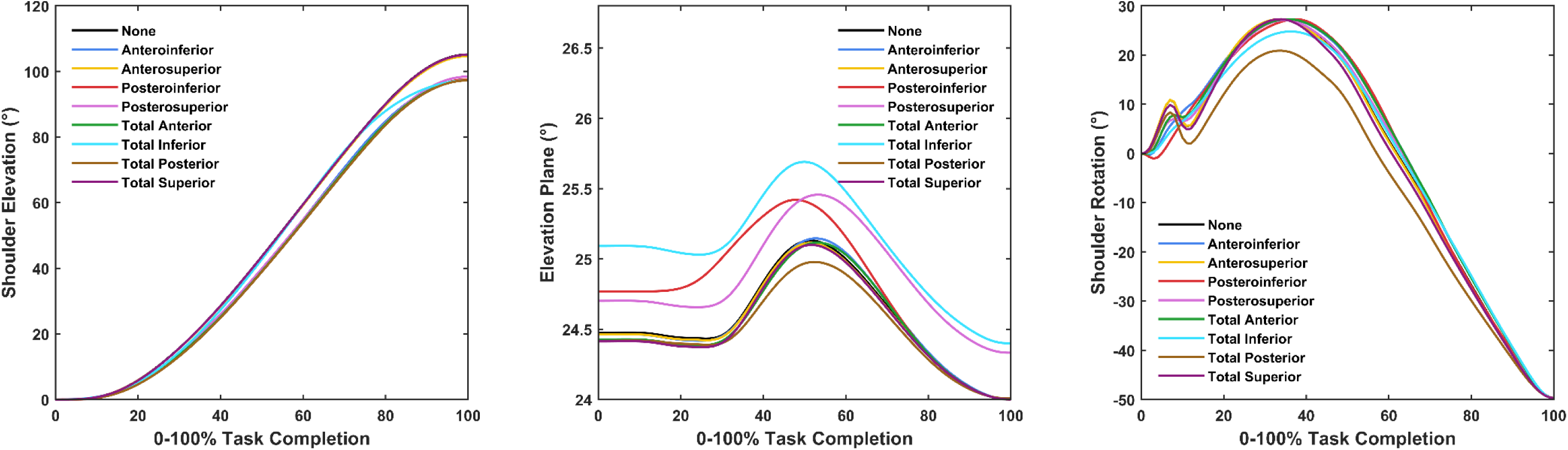
Shoulder kinematics for the baseline (i.e. None) and simulated capsulorrhaphy models during the head touch task. More positive elevation plane values signify elevation shifting further in front of the body (i.e. 90 degree elevation plane refers to pure sagittal plane flexion). Positive and negative shoulder axial rotation values refer to internal and external rotation, respectively.

### Muscle Cost

Across all tasks on average, total muscle cost relative to the ‘None’ condition increased under the selective capsulorrhaphy conditions. The largest increases in relative average total muscle cost came under the total inferior (15.83% ± 16.82%), total anterior (13.19% ± 7.43%), anteroinferior (10.93% ± 11.20%), and total posterior (9.77% ± 13.19%) conditions (see Figure 6A). Inspection of individual tasks revealed variable changes in total muscle cost. An increase in relative total muscle cost was observed across all conditions for the upward reach task – with the greatest increases under the total inferior (20.15%), total anterior (19.74%), total posterior (18.03%) and anteroinferior (15.72%) conditions (see Figure 6B). Total anterior was the only condition to increase relative total muscle cost (5.11%) during the forward reach task – with a decrease observed in all other conditions (see Figure 6C). The changes in relative total muscle cost for the forward reach task were relatively smaller compared to the other two tasks. The largest increases in relative total muscle cost were observed under the total inferior (30.07%), anteroinferior (18.94%), total posterior (16.72%) and total anterior (14.71%) conditions in the head touch task – with the remaining conditions recording minimal change (i.e. < 5%) (see Figure 6D).

**Figure 6.**
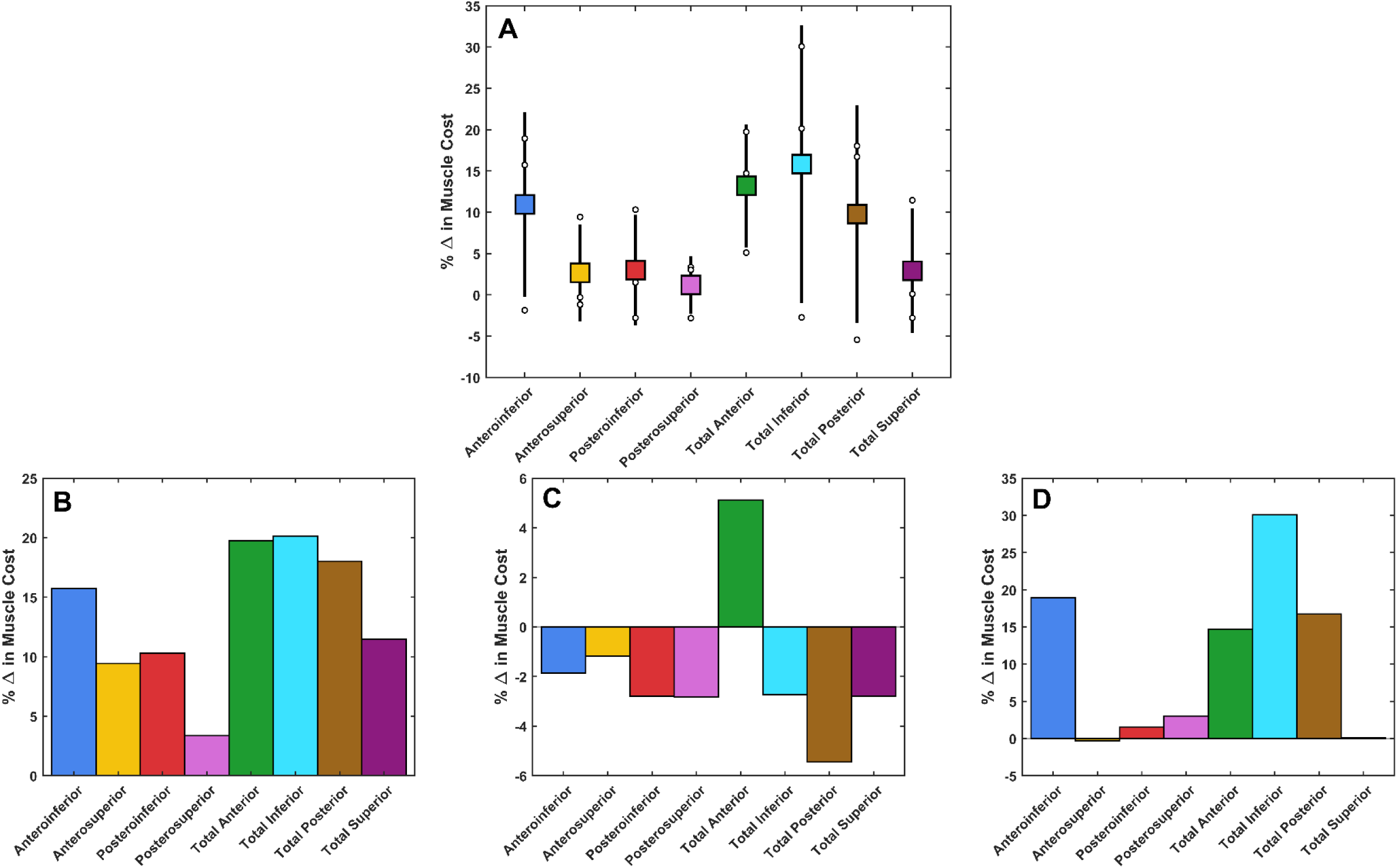
Mean (± standard deviation) percentage change in muscle cost relative to the baseline (i.e. None) model across the three tasks (A). Individual percentage changes in muscle cost for the upward reach (B), forward reach (C) and head touch (D) tasks are also presented.

The anterior deltoid (25.65% ± 3.69% relative contribution to change in muscle cost across simulated capsulorrhaphy conditions), lower trapezius (18.65% ± 7.55%), and lower serratus anterior (15.16% ± 3.92%) were the primary contributors to the increase in total muscle cost during the upward reach task – with smaller contributions made by the upper pectoralis major (6.71% ± 3.98%), supraspinatus (4.63% ± 1.62%), subscapularis (3.48% ± 2.44%) and upper trapezius (3.47% ± 1.05%) (see Supplementary Figure S1). A relatively consistent contribution from a number of muscles appeared to generate the increase in total muscle cost under the total anterior condition during the forward reach task (see Supplementary Figure S2). The lower serratus anterior was the primary contributor to the decrease in total muscle cost under the remaining selective capsulorrhaphy conditions (−19.69% ± 8.80%), with smaller contributions made by the anterior (−7.64% ± 3.13%) and posterior (−6.50% ± 3.52%) deltoids, teres minor (−5.63% ± 7.60%), infraspinatus (−5.55% ± 7.68%) and middle serratus anterior (−4.24% ± 1.33%) (see Supplementary Figure S2). The lower trapezius (13.78% ± 16.24%), anterior deltoid (9.30% ± 11.22%), lower serratus anterior (7.88% ± 12.00%), middle deltoid (7.07% ± 17.63%) and supraspinatus (5.67% ± 3.52%) were the primary contributors to increased total muscle cost during the head touch task – with smaller contributions made by the infraspinatus (3.09% ± 5.39%) and subscapularis (2.46% ± 8.30%) (see Supplementary Figure S3).

### Performance Time

Performance times remained highly consistent (i.e. within 0.05 seconds) across the capsulorrhaphy conditions and all tasks (see Table 4).

**Table 4.** Performance times (in seconds) for selective capsulorrhaphy models across the three simulated tasks

| Capsulorrhaphy Model | Upward Reach | Forward Reach | Head Touch |
| --- | --- | --- | --- |
| None | 0.701 | 0.548 | 0.674 |
| Anteroinferior | 0.653 | 0.548 | 0.622 |
| Anterosuperior | 0.664 | 0.548 | 0.672 |
| Posteroinferior | 0.658 | 0.549 | 0.635 |
| Posterosuperior | 0.674 | 0.548 | 0.642 |
| Total Anterior | 0.648 | 0.547 | 0.621 |
| Total Inferior | 0.657 | 0.549 | 0.645 |
| Total Posterior | 0.656 | 0.548 | 0.629 |
| Total Superior | 0.656 | 0.548 | 0.674 |

## Discussion

The aim of this predictive simulation study was to assess the impact of selective glenohumeral capsulorrhaphy on shoulder kinematics, muscular effort, and performance during relevant functional upper limb movement tasks. Contrary to our hypotheses, shoulder joint kinematics and task performance remained relatively stable during the simulated tasks. An overall trend for increased muscular effort (typical range of 5-30% increase) under the selective capsulorrhaphy conditions was observed, however this response was dependent on the task and selective capsulorrhaphy condition.

The simulated selective capsulorrhaphy conditions had a minimal impact on the movement strategies used and task performance time. While kinematics and performance time did fluctuate across the different models, the majority of changes were relatively small (i.e. 1-2 degrees and 0.05 seconds). These findings suggest that with the increases in muscle activation and forces observed, the performance output of the functional tasks assessed in this study can be maintained. It is important to note that the present study simulated the average reduction in range of motion following selective capsulorrhaphy^1^ and the subsequent effect of this on movement and muscle function. Both larger and smaller effects on range of motion were observed across the individual cadaveric specimen by Gerber et al.^1^ – with quite large ranges (e.g. over 50°) observed between the worst- and best-case scenarios in certain joint positions. It may be that greater changes in movement strategies and task performance would present under the scenarios where passive range of motion is severely hampered. The models and code provided with this paper (https://simtk.org/projects/gh-caps-sims) can be used for replication experiments and provide an opportunity to assess the impact of larger or smaller reductions in passive range of motion.

Certain capsulorrhaphy conditions induced slightly larger changes in shoulder joint kinematics during specific tasks. First, a small reduction in shoulder elevation was observed during the head touch task under the anteroinferior, posteroinferior, posterosuperior, total anterior, total inferior and total posterior conditions – with this being most evident towards the end of the task at the highest degrees of elevation. Second, the total posterior condition largely varied from the remaining conditions for the degree of shoulder internal rotation during the head touch task – demonstrating reduced internal rotation, particularly during the first half of the task. These results are not surprising when considering the specific impacts on range of motion these capsulorrhaphy conditions presented. Anteroinferior, posteroinferior, posterosuperior, total anterior, total inferior and total posterior conditions generate the largest reductions for elevation range of motion in abduction.^1^ Similarly, the total posterior condition incurs the largest reductions in shoulder internal rotation range of motion at lower degrees of shoulder elevation (i.e. between zero and 45°).^1^ Subsequently – larger passive resistive forces were experienced by the model in the present study under these capsulorrhaphy conditions when generating abduction and internal rotation movements at lower elevation angles, respectively. The head touch task was performed at a lower elevation plane angle compared to the reaching tasks – signifying the arm was raised more so in the frontal versus sagittal plane. This is likely why shoulder elevation and internal rotation were more readily altered under the aforementioned capsulorrhaphy conditions during the head touch, but not other, tasks.

In general, muscle cost increased under the selective capsulorrhaphy conditions when compared to the ‘None’ model. Compared to the ‘None’ model, all capsulorrhaphy conditions involved an increase in the passive resistance to shoulder motion (see Supplementary Figures S10–S13). It is therefore logical that under these conditions the muscles were required to increase their activation to produce more force to counter the additional passive resistance – and hence an increase in cost would be observed. This was the case in the present study, whereby greater activation and forces were observed in the specific muscles responsible for elevating muscle cost (see Supplementary Figures S4–S9). This behaviour was predominantly observed in the prime movers for each task – such as the anterior and middle deltoids for the overhead (i.e. upward reach and head touch) tasks – and was the major factor in increasing muscle cost. The minimal changes in joint kinematics meant that muscles remained operating at similar lengths relative to optimal. Similarly, we did not observe a dramatic change in muscle contraction velocity across the simulated capsulorrhaphy conditions.

The increase in total muscle cost under the majority of capsulorrhaphy conditions highlights an increase in load on the muscles, which could lead to fatigue or muscle damage.^10^ This may present a significant problem for individuals who more readily repeat the functional tasks we simulated in their daily activities (e.g. manual handling workers), or for those who perform tasks with large degrees of shoulder movement that near full range of motion (e.g. overhead lifting tasks). The presence of elevated muscle loading early on in the recovery from surgery could also present a risk to the joint capsule repair. Inadequate muscle function stemming from fatigue or damage may not maintain the glenohumeral joint centre of rotation during movement, risking instability and damage to the surgically repaired capsule, or overload the rotator cuff tendons.^31,32^ Rehabilitation following glenohumeral capsulorrhaphy must consider these factors, and likely include a focus on strengthening the muscles responsible for driving movements commonly performed by the individual. It may also be advisable to avoid shoulder postures near full range of motion that could significantly elevate muscle loads, particularly early on in recovery when some muscle weakness may be present.

Changes in muscle cost were relatively smaller for the forward reach task compared to the upward reach and head touch tasks. This finding is likely due to the larger joint angle ranges achieved during the upward reach and head touch tasks. Shoulder elevation and rotation during the forward reach peaked at approximately 40 and 24 degrees, respectively. In contrast, larger peaks were observed for shoulder elevation (i.e. 90-100 degrees) and rotation (i.e. 30-40 degrees) in the other tasks. The passive structures at the glenohumeral joint (i.e. ligaments, joint capsule) are responsible for restricting glenohumeral motion at end-, rather than mid-range humeral elevation.^33^ Our findings appear to support this notion, whereby the model structures simulating passive restraints produced more resistive force closer to end-range shoulder elevation – and the largest effects were therefore observed in tasks where shoulder elevation was closer to this point (i.e. ∼90-100 degrees). While an increase in passive resistance would still be experienced at lower degrees of shoulder elevation, the magnitude of this did not appear large enough to generate substantial (or any) increases in total muscle cost. In conjunction with targeted rehabilitation therapy, individuals who have undergone glenohumeral capsulorrhaphy may be able to minimise the stress placed on muscles by limiting activities that involve high degrees of shoulder elevation. This is, however, only a temporary solution as patients will likely need to return to full range of motion in order to function in routine daily activities. Standard rehabilitation strategies following shoulder stabilisation that focus on muscle strength and control^34,35^ are supported by our results. In particular, strengthening the major shoulder muscles will serve to counter the additional passive resistance and potential increase in muscle cost incurred by the glenohumeral capsulorrhaphy.

Total anterior capsulorrhaphy was the only simulated condition to increase muscle cost relative to the ‘None’ model across all tasks. All of the simulated movement tasks in the present study were performed using shoulder flexion (i.e. elevation angle > 0) and with internal axial rotation. The total anterior capsulorrhaphy often induced the largest reductions in shoulder flexion and internal rotation range of motion,^1^ and hence additional resistance to these motions was present under this condition. The anteroinferior, and total anterior, inferior and posterior conditions resulted in the largest increases in total muscle cost in the upward reach and head touch tasks. Again, these capsulorrhaphy conditions typically demonstrated the largest impacts on range of motion loss in both shoulder elevation and internal rotation.^1^ The findings of our study appear to support a relationship between the range of motion deficit induced by the glenohumeral capsulorrhaphy and an increase in muscle cost. We propose that individuals who experience substantial losses in range of motion following glenohumeral capsulorrhaphy may be at greater risk of musculoskeletal issues (i.e. muscle fatigue and damage, joint instability, and tendon loading) induced by increased muscle loads. Certain selective capsulorrhaphy conditions also have a prominent effect on limiting external rotation (e.g. total superior) and abduction (e.g. total inferior) range of motion.^1^ While we did not test any movements that included abduction and external rotation actions, our results indicate that increases in muscle cost would also be observed in such tasks – with these increases likely higher under the capsulorrhaphy conditions that have a greater impact on range of motion in these directions.

The present study demonstrates how musculoskeletal modelling and predictive simulation can be used to assess potential changes in neuromuscular function and movement during upper limb tasks following a specific adaptation in the musculoskeletal system (i.e. increased passive resistance following glenohumeral capsulorrhaphy). Our study is limited in that it employed a generic musculoskeletal model, literature-based values for the reductions in shoulder range of motion, and simulated task performance based on a generic set of goals. A similar approach could, however, be taken in a patient-specific manner, whereby – (i) a patient-specific model of the upper limb (e.g. developed from medical imaging data); (ii) patient-specific ranges of motion, and expected or measured losses in range of motion following surgery; and (iii) kinematic data experimentally measured from the patient (e.g. via motion capture, video analysis or wearable sensors) during functional tasks relative to their daily performance – could be used. Such an approach may be useful in providing a patient-specific understanding of how an already performed or planned glenohumeral capsulorrhaphy procedure may impact the individual’s upper limb function. This information could then be used to design a post-surgery rehabilitation plan that targeted the specific muscles that may see an increased load in the individual. Embedding patient-specific musculoskeletal modelling and simulation procedures within clinical practice could provide added value across a wide spectrum of patients. However, efforts must be made to ensure this type of analysis and planning is easily accessible to clinicians and can be efficiently integrated into clinical practice.

There are certain limitations relating to the modelling approaches used in our study that need to be considered when interpreting the results. First, we chose to lock motion of the trunk to ensure that the desired movement goal was achieved only by the upper limb under the different capsulorrhaphy conditions. This was necessary to ensure that adaptations to counter the additional passive resistance were achieved by the shoulder muscles – however, it may not represent certain adaptive movement strategies an individual uses following glenohumeral capsulorrhaphy. In the face of added passive resistance at the shoulder, an individual may choose to use the trunk to a greater extent (i.e. sagittal or frontal plane leaning) to achieve reaching or overhead movements. This type of behaviour has been observed in individuals with shoulder pain during a fatiguing reaching task – whereby those with pain tend to move their centre of mass more (i.e. greater trunk involvement) to perform the task.^36^ Greater trunk involvement would likely reduce muscular involvement at the shoulder, subsequently reducing muscle cost, but could increase the moment arm of the trunk relative to the spinal joints. Therefore, there is a likely trade-off in potentially shifting load from the shoulder to the spine with such an approach. Future predictive simulations would need to consider appropriate cost functions to limit or balance the involvement of the trunk during upper limb movements. Second, we did not include a constraint on the resultant glenohumeral joint reaction vector as has been done in previous studies.^20,37–39^ Existing work has constrained the resultant glenohumeral joint reaction vector so that its projection remains within an ellipse representing the glenoid face,^37,38^ or ensured that the glenohumeral compressive force outweighs the shear components.^20,39^ The goal of these constraints is to ensure the simulated muscle activity includes appropriate rotator cuff function to maintain glenohumeral joint stability.^20,37–39^ Rotator cuff activation and force may therefore have been underestimated in our study. However, this remained consistent across the different simulated capsulorrhaphy conditions, and hence we believe our estimates of increased muscle cost relative to the ‘None’ condition are still appropriate. Implementing such a constraint within the current OpenSim Moco framework is also difficult and would likely result in a substantial increase in simulation times. Future predictive simulation studies, particularly those with an interest in glenohumeral (in)stability and/or rotator cuff function (including post joint replacement surgery), should still consider this inclusion. Third, we only used a single participant design with an ‘optimised’ movement strategy. Different effects on shoulder kinematics and muscle function or cost may be observed under different baseline movement strategies. For example, an individual who performs the functional tasks differently to what we observed in our ‘None’ condition may need to adapt their kinematic strategy in a different manner to adjust to the additional passive resistance induced by a glenohumeral capsulorrhaphy procedure. Future predictive simulation studies could include participant-specific movement strategies as a baseline condition and observe how a model adapts to added passive resistance under these initial conditions.

## Conclusions

Our predictive simulation study found that shoulder kinematics and performance times did not dramatically change during simulated upper limb movement tasks under various simulated capsulorrhaphy conditions. Despite the lack of kinematic and performance changes, glenohumeral capsulorrhaphy generally resulted in an increase in total muscle cost during tasks. Larger increases in muscle cost were typically observed under the simulated total inferior, total anterior, anteroinferior and total posterior conditions. Changes in muscle cost were, however, dependent on the task being performed. The largest increases in muscle cost were observed in tasks that required shoulder kinematics closer to end range of motion. Elevated muscle loading, particularly early on in recovery, could present a risk to the joint capsule repair. Our results highlight the need for appropriate guided and targeted rehabilitation following glenohumeral capsulorrhaphy to account for the elevated demands placed on the muscle, particularly when significant range of motion loss presents.

## Supporting information

Supplementary Document 1 - Grid Density Refinement

## Notes

https://simtk.org/projects/gh-caps-sims

### Competing Interest Statement

The authors have declared no competing interest.

### Summary of Updates

Updated documents to include doi

