## Supplementary Document 1 - Grid Density Refinement for "Simulating the Impact of Glenohumeral Capsulorrhaphy on Movement Kinematics and Muscle Function in Activities of Daily Living"

### A complete summary of grid refinement simulations for assessing an appropriate node density for each simulated task

All node densities (i.e. 25, 51, 101, 151, 201 and 251 nodes) for all tasks converged to an optimal solution and satisfied the task constraints. The objective function value and task performance time remained relatively consistent across node densities for all tasks – suggesting finer grids did not drastically change overall task performance (see Figure SD1).

**A)** **B)**

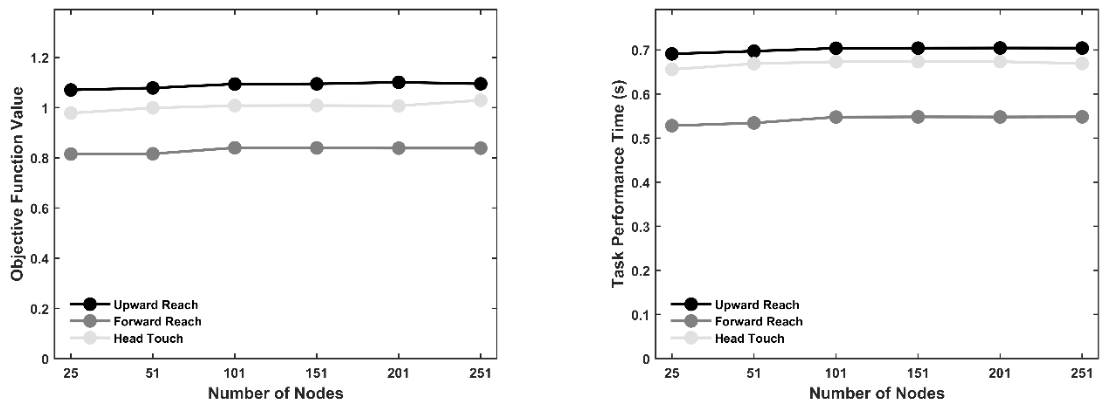


**Figure SD1** **Objective function (A) and performance time (B) values for the simulated tasks across different grid densities.**

#### *Upward Reach Task*

The root mean square error between adjacent node solutions plateaued at 201 nodes for joint angles and at 151 nodes for muscle activations (see Figure SD8). There was little visual difference in the joint angles and muscle activations from 151 nodes and up (see Figures SD2 and SD3, respectively) – with no dramatic increase in solver duration (see Figure SD8). Based on these data, a node density of 201 was selected for the upward reach task.

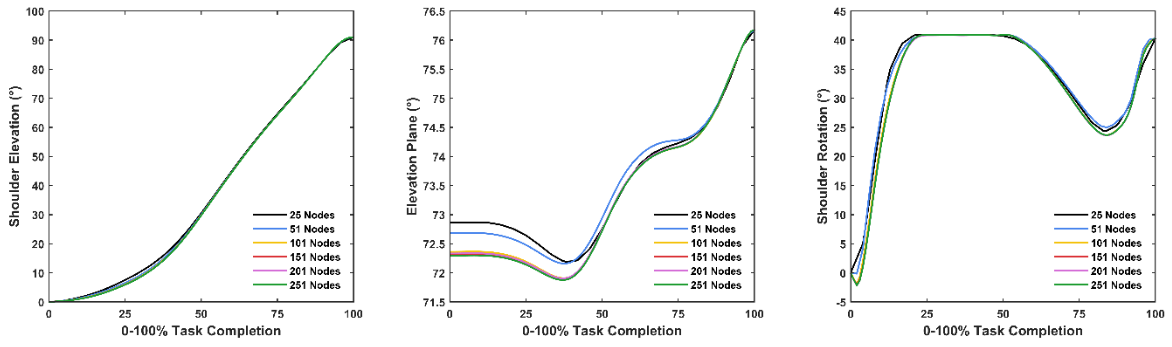


**Figure SD2** **Comparison of time-normalised shoulder joint angles for the upward reach task across different grid densities. More positive elevation plane values signify elevation shifting further in front of the body (i.e. 90 degree elevation plane refers to pure sagittal plane flexion). Positive and negative shoulder axial rotation values refer to internal and external rotation, respectively.**

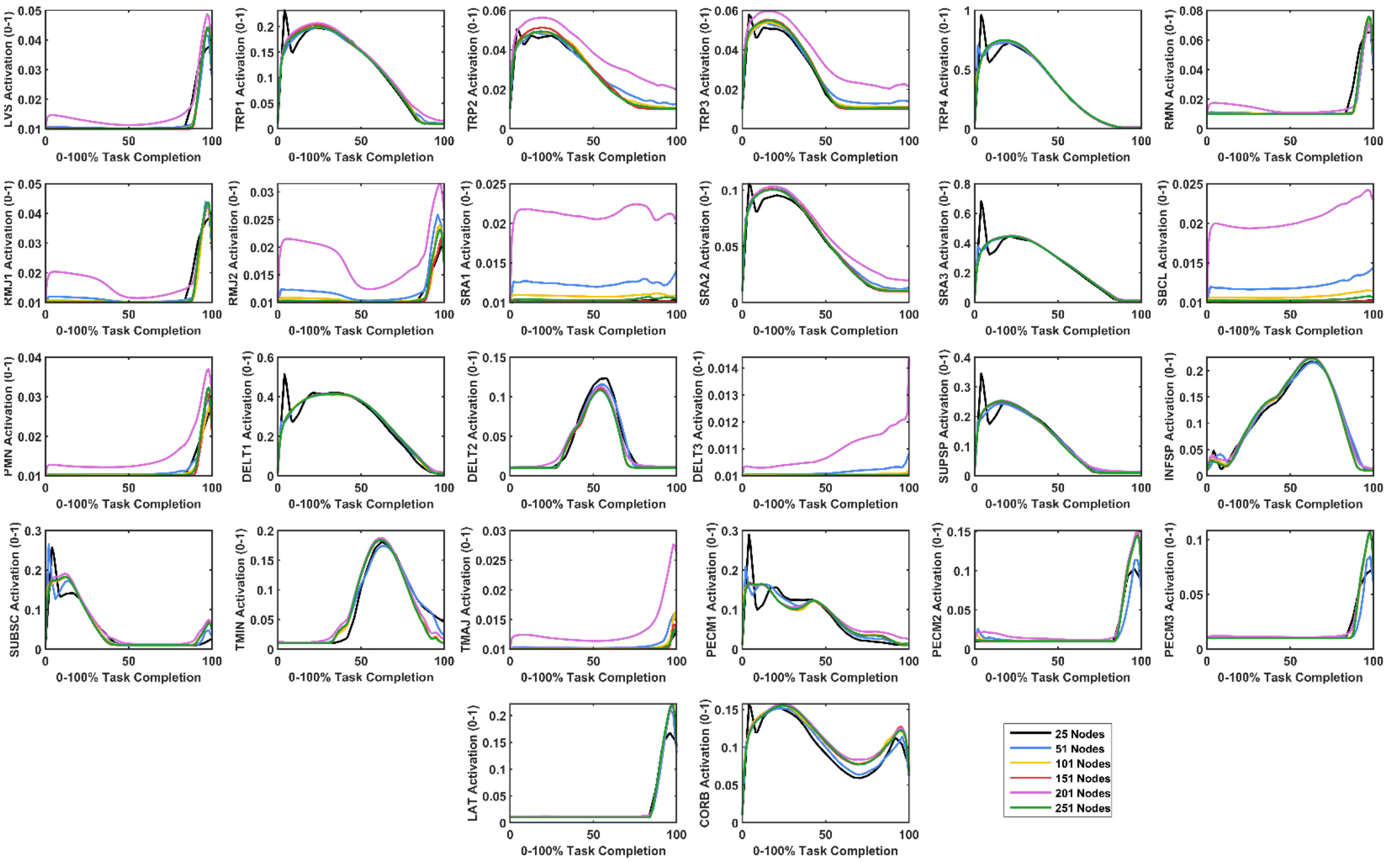


**Figure SD3** **Comparison of time-normalised muscle activations for the upward reach task across different grid densities. LVS – levator scapulae; TRP1 – superior trapezius; TRP2 – upper-middle trapezius; TRP3 – lower-middle trapezius; TRP4 – lower trapezius; RMN – rhomboid minor; RMJ1 – upper rhomboid major; RMJ2 – lower rhomboid major; SRA1 – upper serratus anterior; SRA2 – middle serratus anterior; SRA3 – lower serratus anterior; SBCL – subclavius; PMN – pectoralis minor; DELT1 – anterior deltoid; DELT2 – middle deltoid; DELT3 – posterior deltoid; SUPSP – supraspinatus; INFSP – infraspinatus; SUBSC – subscapularis; TMIN – teres minor; TMAJ – teres major; PECM1 – upper pectoralis major; PECM2 – middle pectoralis major; PECM3 – lower pectoralis major; LAT – latissimus dorsi; CORB – coracobrachialis.**

### *Forward Reach Task*

The root mean square error between adjacent node solutions plateaued at 201 nodes for joint angles and at 101 nodes for muscle activations (see Figure SD8). There was little visual difference in the joint angles and muscle activations from 101 nodes and up (see Figures SD4 and SD5, respectively) – with no dramatic increase in solver duration (see Figure SD8). Based on these data, a node density of 201 was selected for the forward reach task.

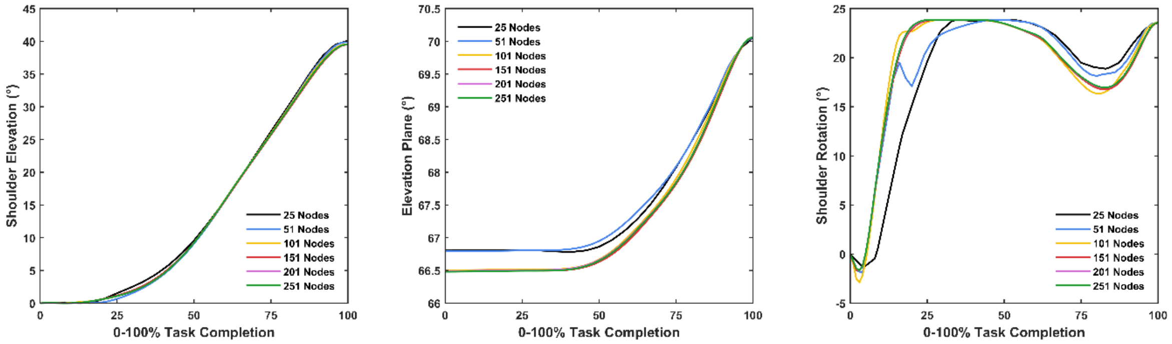


**Figure SD4** **Comparison of time-normalised shoulder joint angles for the forward reach task across different grid densities. More positive elevation plane values signify elevation shifting further in front of the body (i.e. 90 degree elevation plane refers to pure sagittal plane flexion). Positive and negative shoulder axial rotation values refer to internal and external rotation, respectively.**

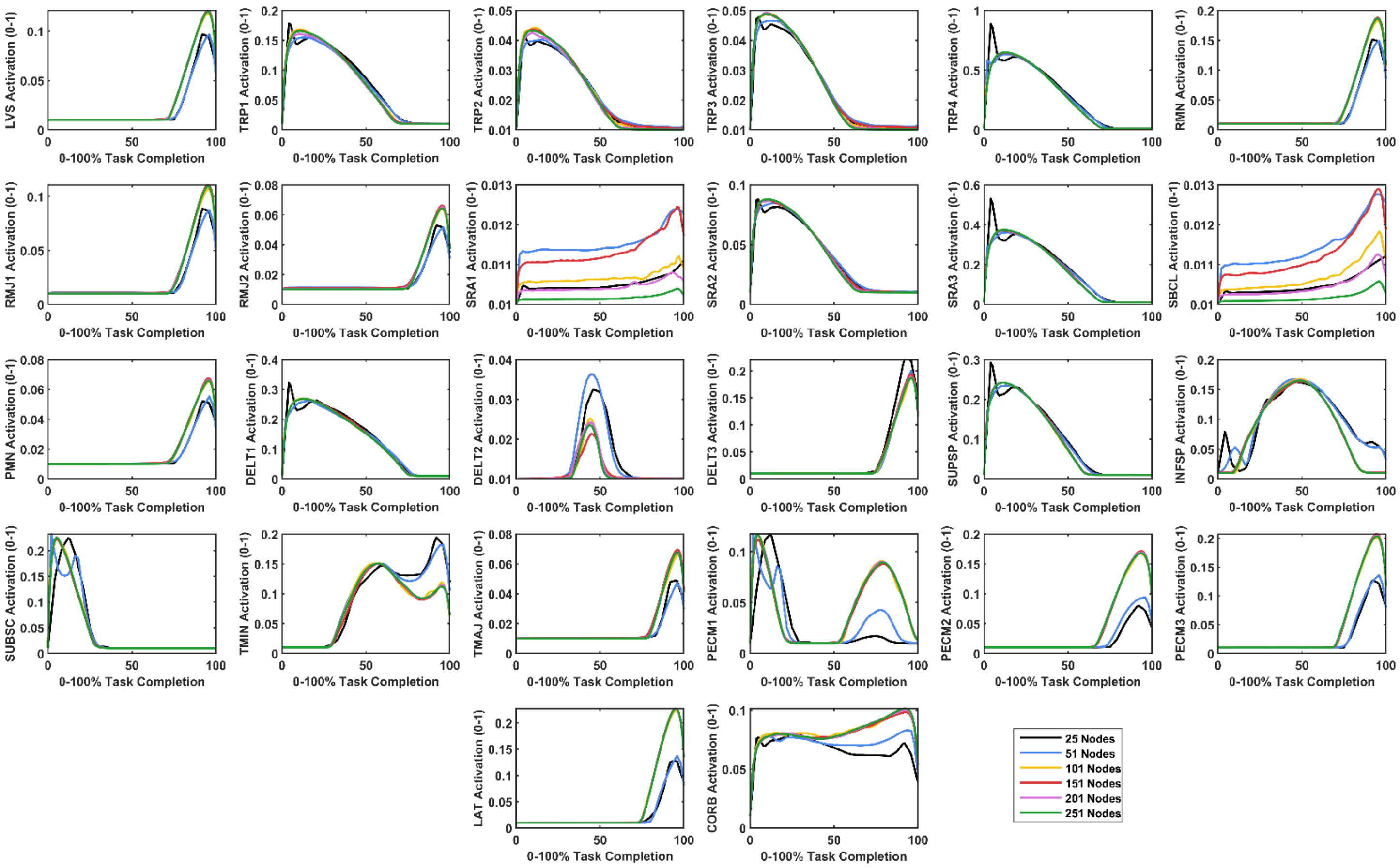


**Figure SD5** **Comparison of time-normalised muscle activations for the forward reach task across different grid densities. LVS – levator scapulae; TRP1 – superior trapezius; TRP2 – upper-middle trapezius; TRP3 – lower-middle trapezius; TRP4 – lower trapezius; RMN – rhomboid minor; RMJ1 – upper rhomboid major; RMJ2 – lower rhomboid major; SRA1 – upper serratus anterior; SRA2 – middle serratus anterior; SRA3 – lower serratus anterior; SBCL – subclavius; PMN – pectoralis minor; DELT1 – anterior deltoid; DELT2 – middle deltoid; DELT3 – posterior deltoid; SUPSP – supraspinatus; INFSP – infraspinatus; SUBSC – subscapularis; TMIN – teres minor; TMAJ – teres major; PECM1 – upper pectoralis major; PECM2 – middle pectoralis major; PECM3 – lower pectoralis major; LAT – latissimus dorsi; CORB – coracobrachialis.**

### *Head Touch Task*

The root mean square error between adjacent node solutions plateaued at 101 nodes for joint angles and at 151 nodes for muscle activations (see Figure SD8). Root mean square error increased for the head touch task across the various measures at 201 or 251 nodes (see Figure SD8). There was little visual difference in the joint angles and muscle activations from 101 to 201 nodes (see Figures SD6 and SD7, respectively) – with no dramatic increase in solver duration (see Figure SD8). The 251 node solution induced noisy muscle activation signals with a substantial increase in solver duration. Based on these data, a node density of 151 was selected for the head touch task.

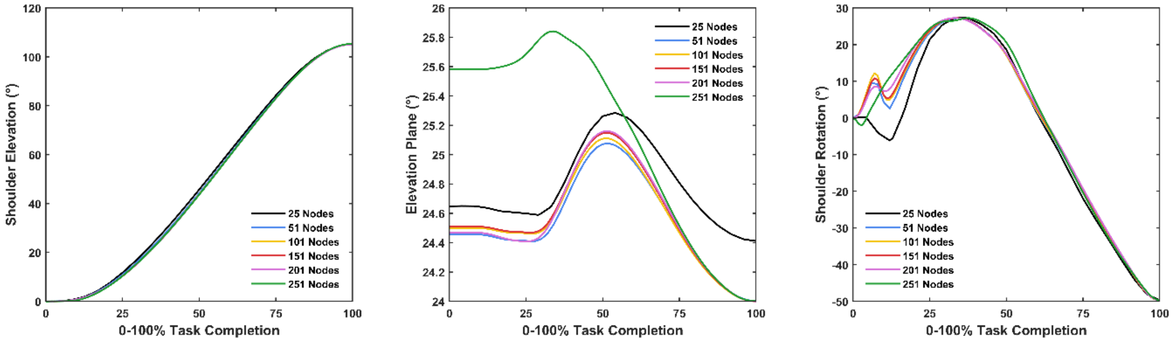


**Figure SD6** **Comparison of time-normalised shoulder joint angles for the head touch task across different grid densities. More positive elevation plane values signify elevation shifting further in front of the body (i.e. 90 degree elevation plane refers to pure sagittal plane flexion). Positive and negative shoulder axial rotation values refer to internal and external rotation, respectively.**

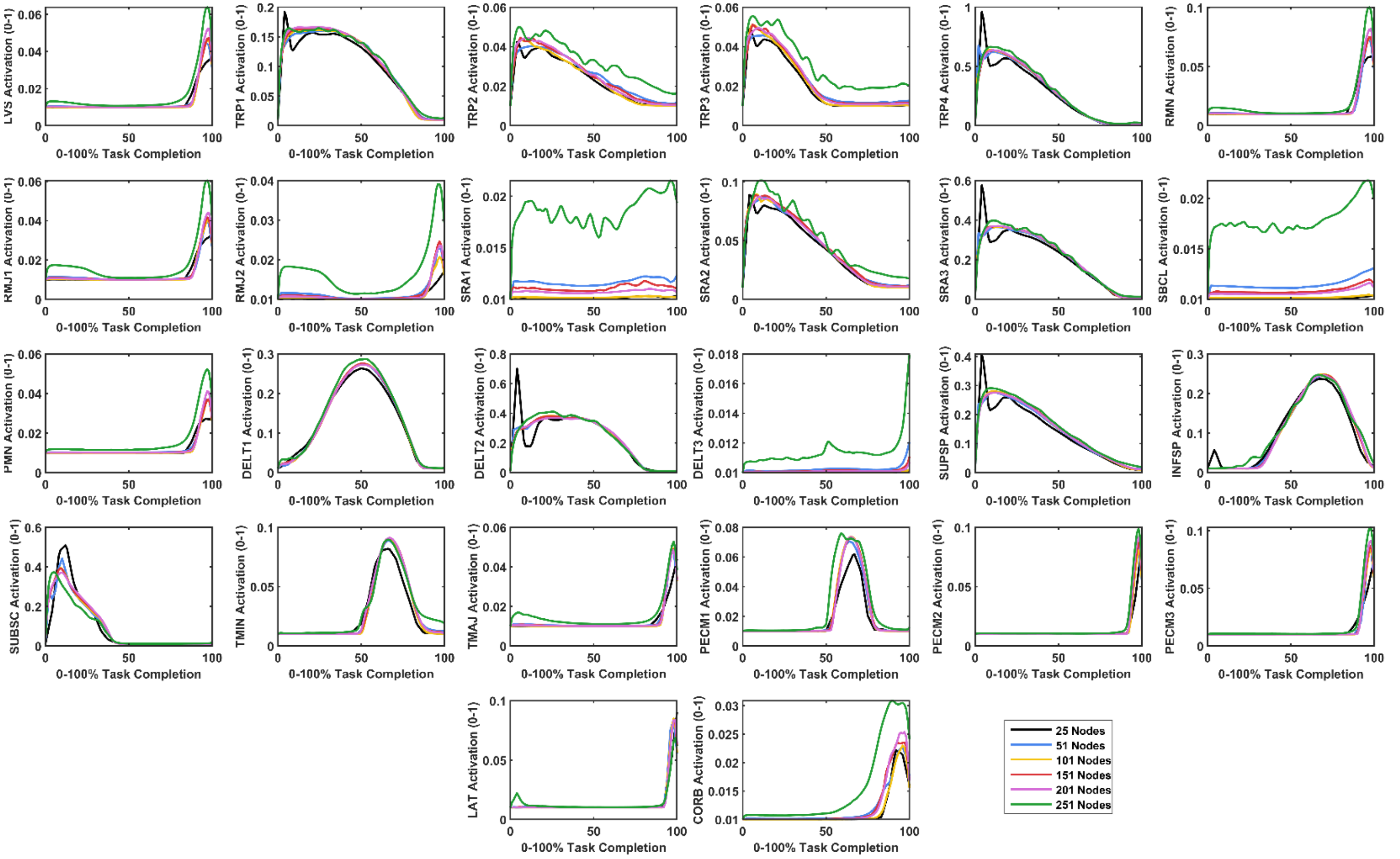


**Figure SD7** **Comparison of time-normalised muscle activations for the head touch task across different grid densities. LVS – levator scapulae; TRP1 – superior trapezius; TRP2 – upper-middle trapezius; TRP3 – lower-middle trapezius; TRP4 – lower trapezius; RMN – rhomboid minor; RMJ1 – upper rhomboid major; RMJ2 – lower rhomboid major; SRA1 – upper serratus anterior; SRA2 – middle serratus anterior; SRA3 – lower serratus anterior; SBCL – subclavius; PMN – pectoralis minor; DELT1 – anterior deltoid; DELT2 – middle deltoid; DELT3 – posterior deltoid; SUPSP – supraspinatus; INFSP – infraspinatus; SUBSC – subscapularis; TMIN – teres minor; TMAJ – teres major; PECM1 – upper pectoralis major; PECM2 – middle pectoralis major; PECM3 – lower pectoralis major; LAT – latissimus dorsi; CORB – coracobrachialis.**

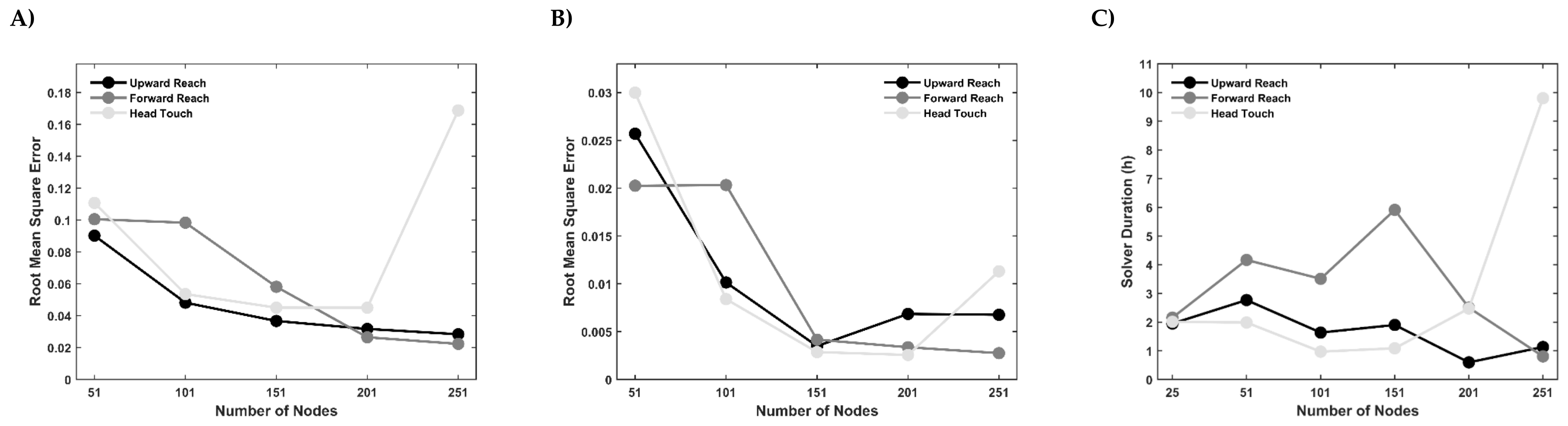


**Figure SD8** **Root mean square error (RMSE) values as a result of increasing grid density for shoulder joint angles (A) and muscle activations (B), and solver duration for the simulated tasks across different grid densities (C).**
