## Supplementary Figures for "Simulating the Impact of Glenohumeral Capsulorrhaphy on Movement Kinematics and Muscle Function in Activities of Daily Living"

The following figures present:

1) The relative contribution of individual muscles to the absolute change in muscle cost across the three simulated tasks under the various simulated glenohumeral capsulorrhaphy conditions.
2) Normalised activations of individual muscles across the three simulated tasks under the various simulated glenohumeral capsulorrhaphy conditions.
3) The forces produced by individual muscles across the three simulated tasks under the various simulated glenohumeral capsulorrhaphy conditions.
4) The resistive torques generated by the simulated passive restraints across the three simulated tasks under the various simulated glenohumeral capsulorrhaphy conditions.

**Abbreviations**

LVS – levator scapulae

SUBCL – subclavius

TRP1 – superior trapezius

TRP2 – upper-middle trapezius

TRP3 – lower-middle trapezius

TRP4 – lower trapezius

RMN – rhomboid minor

RMJ1 – superior rhomboid major

RMJ2 – lower rhomboid major

SRA1 – superior serratus anterior

SRA2 – middle serratus anterior

SRA3 – lower serratus anterior

CORB – coracobrachialis

DELT1 – anterior deltoid

DELT2 – middle deltoid

DELT3 – posterior deltoid

SUBSC – subscapularis

SUPSP – supraspinatus

INFSP – infraspinatus

LAT – latissimus dorsi

TMIN – teres minor

TMAJ – teres major

PMN – pectoralis minor

PECM1 – superior pectoralis major

PECM2 – middle pectoralis major

PECM3 – lower pectoralis major

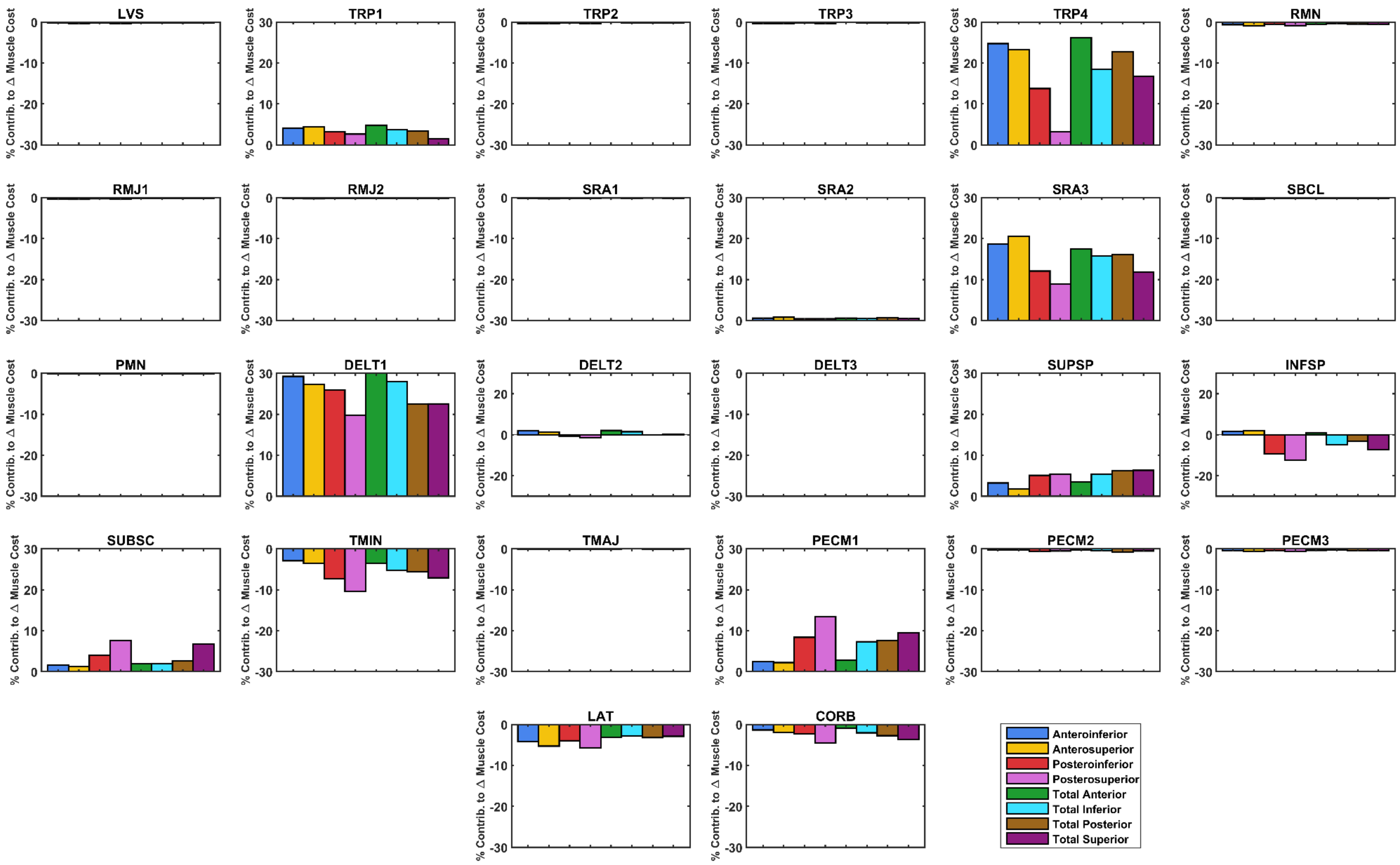


**Figure S1** **The relative contribution of individual muscles to the absolute change in total muscle cost for the upward reach task.**

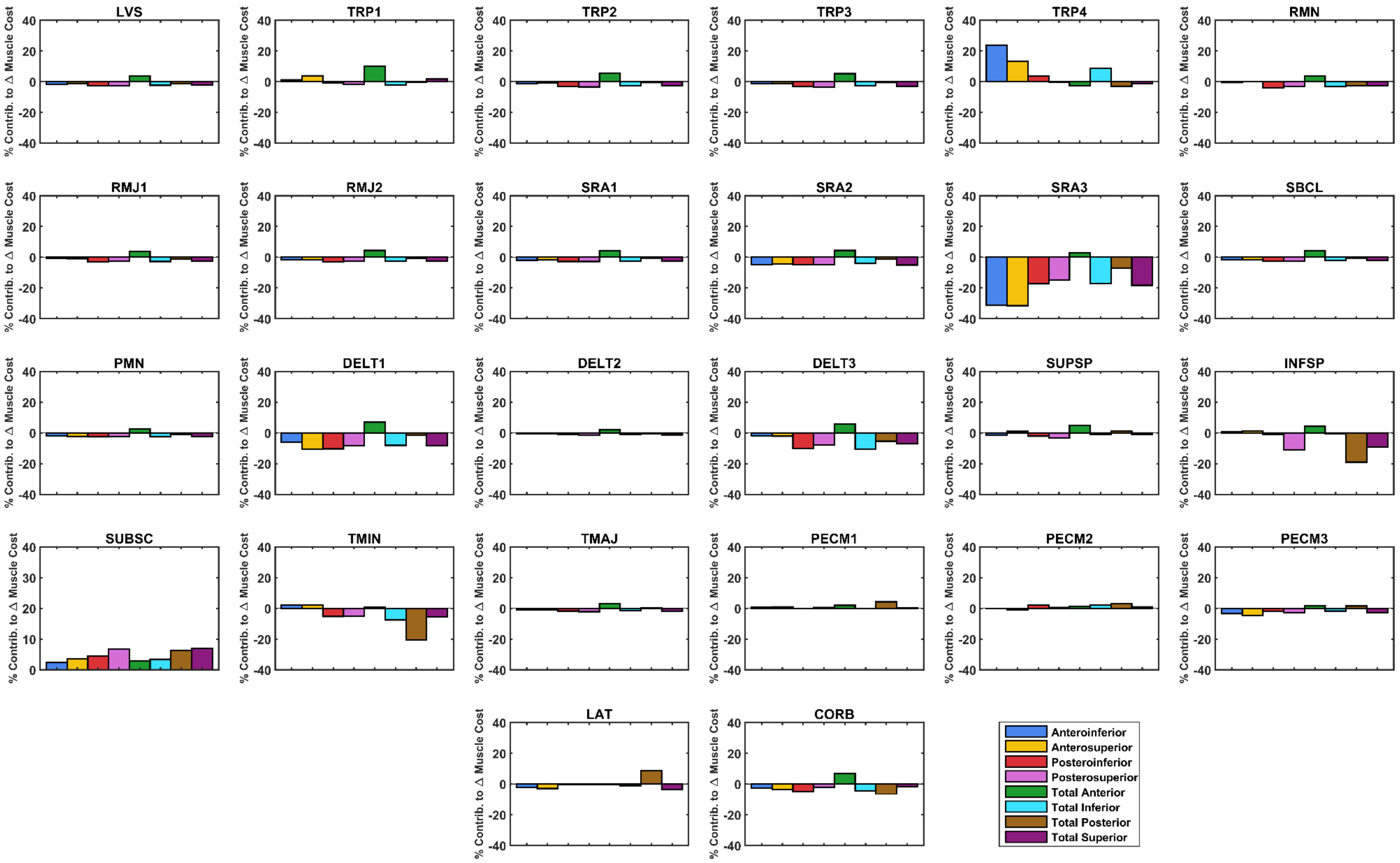


**Figure S2** **The relative contribution of individual muscles to the absolute change in total muscle cost for the forward reach task.**

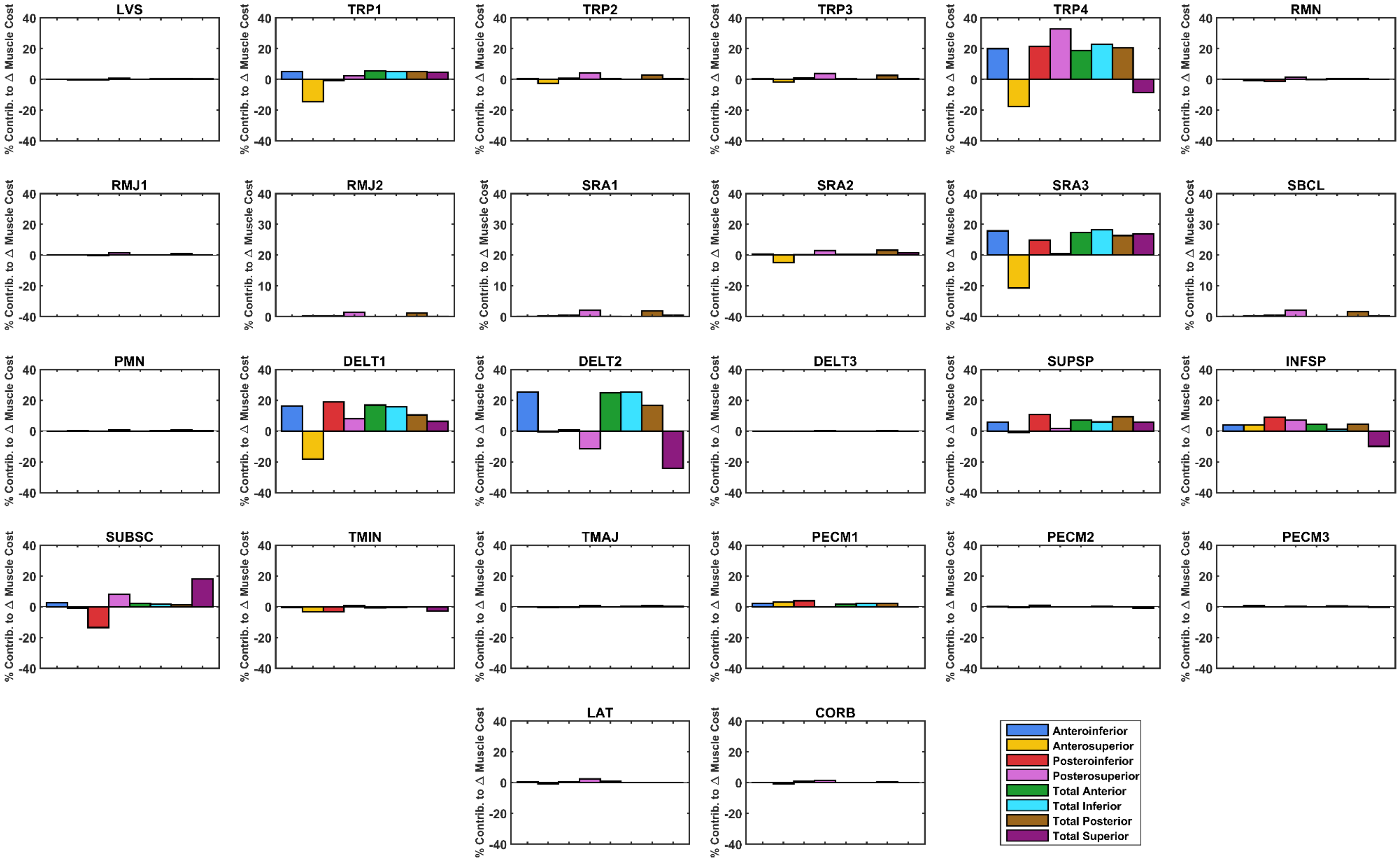


**Figure S3** **The relative contribution of individual muscles to the absolute change in total muscle cost for the head touch task.**

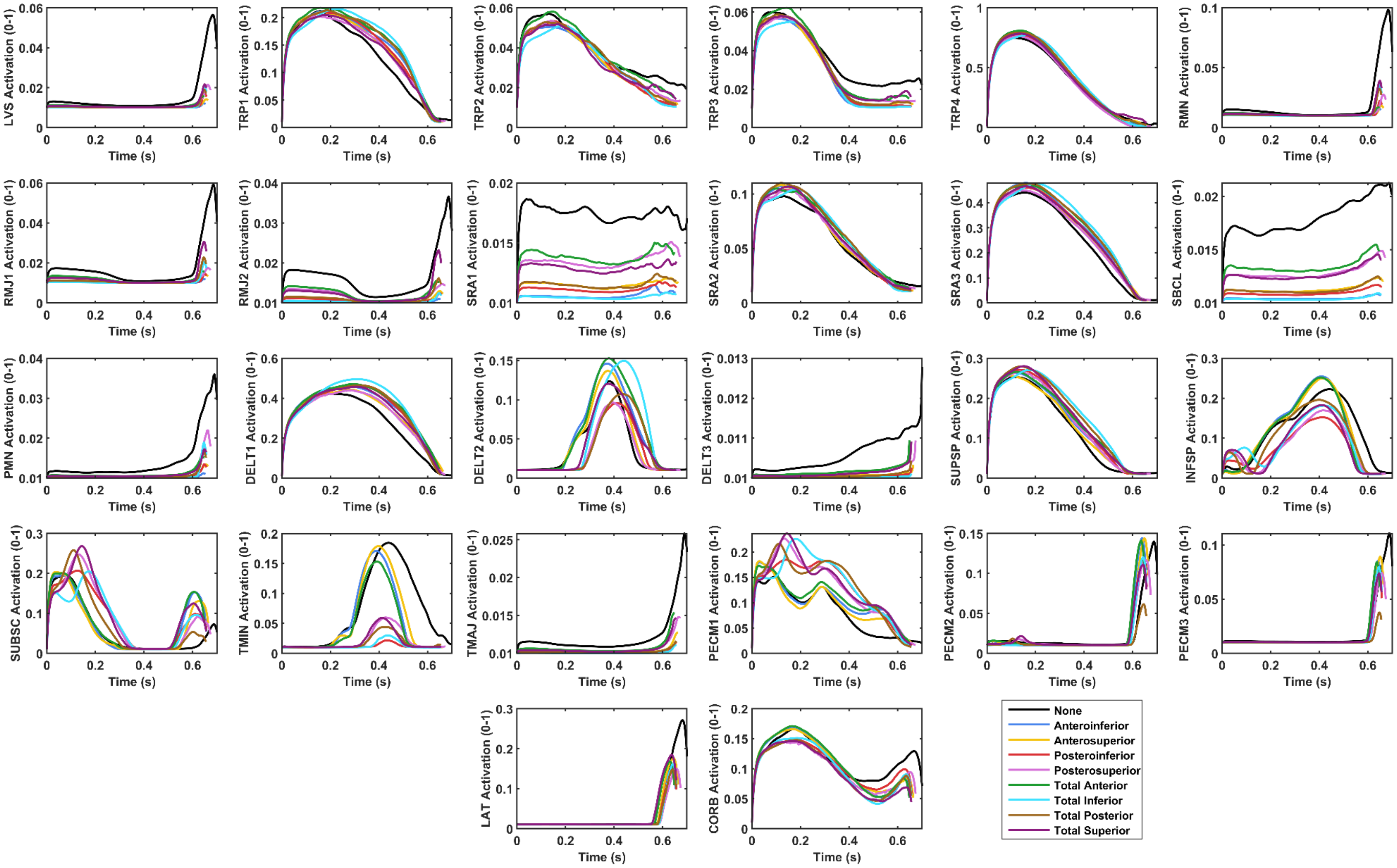


**Figure S4** **Muscle activations (0 – 1 of maximal activation) of individual muscles for the upward reach task.**

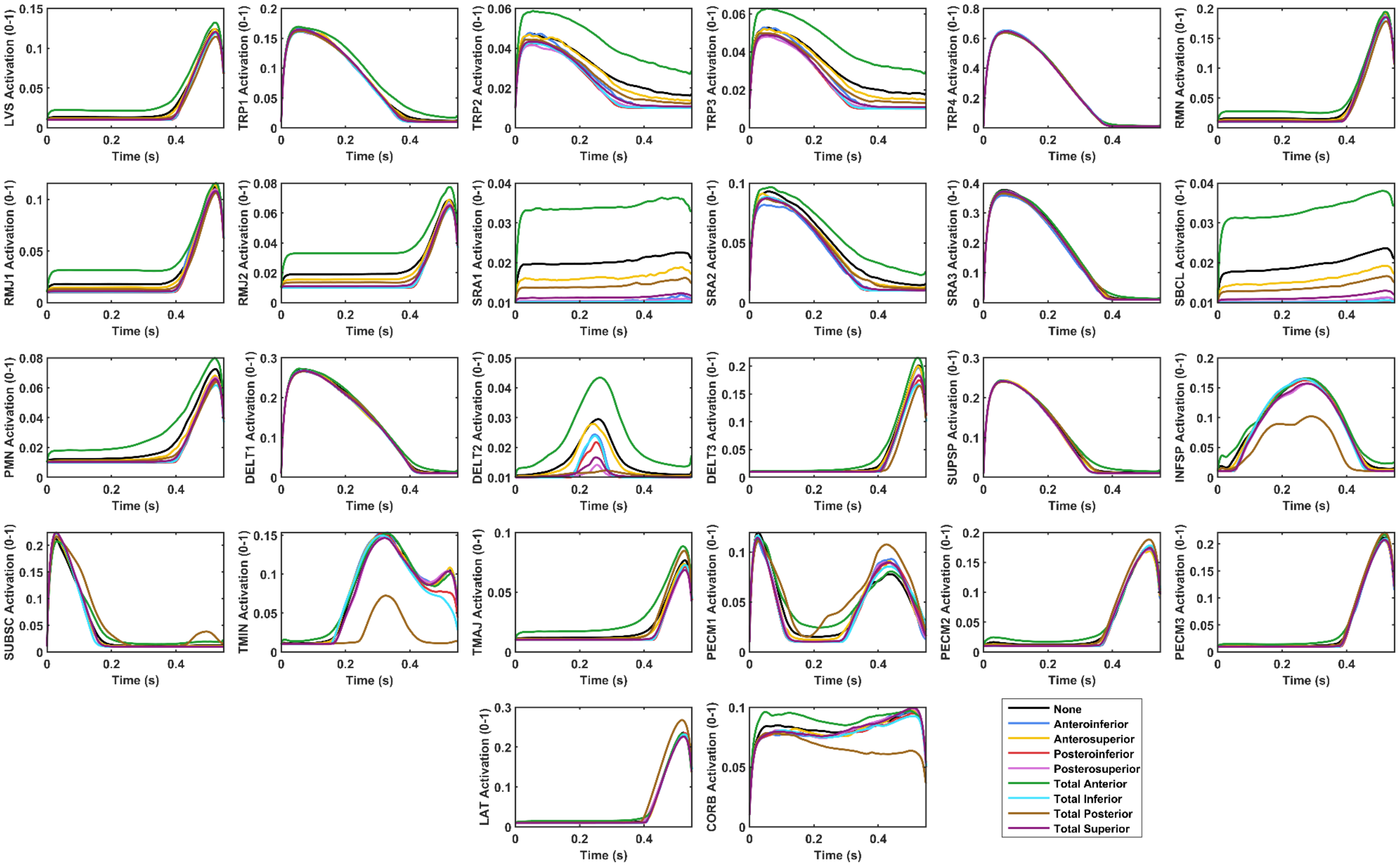


**Figure S5** **Muscle activations (0 – 1 of maximal activation) of individual muscles for the forward reach task.**

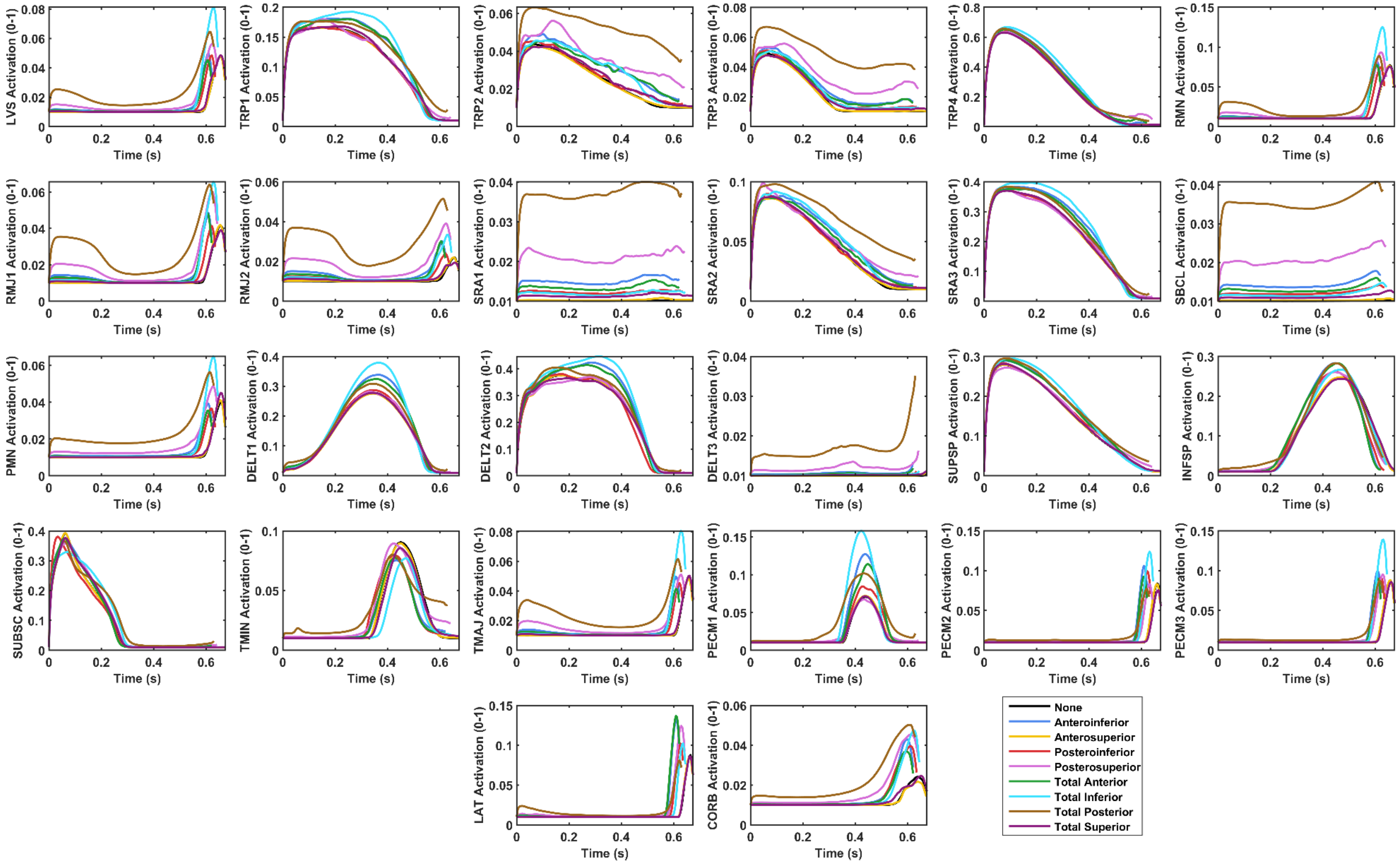


**Figure S6** **Muscle activations (0 – 1 of maximal activation) of individual muscles for the head touch task.**

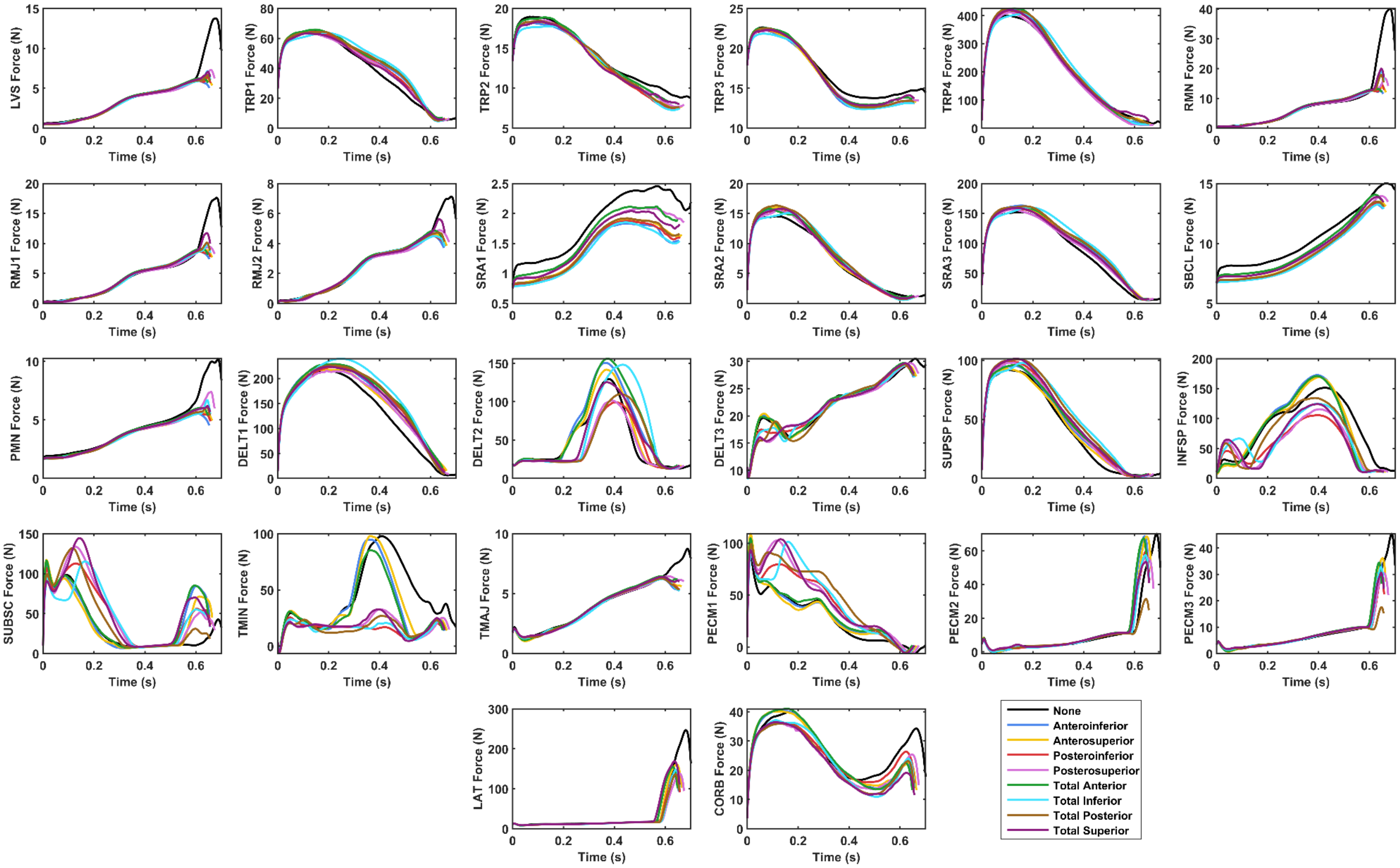


**Figure S7** **Muscle forces (in Newtons) of individual muscles for the upward reach task.**

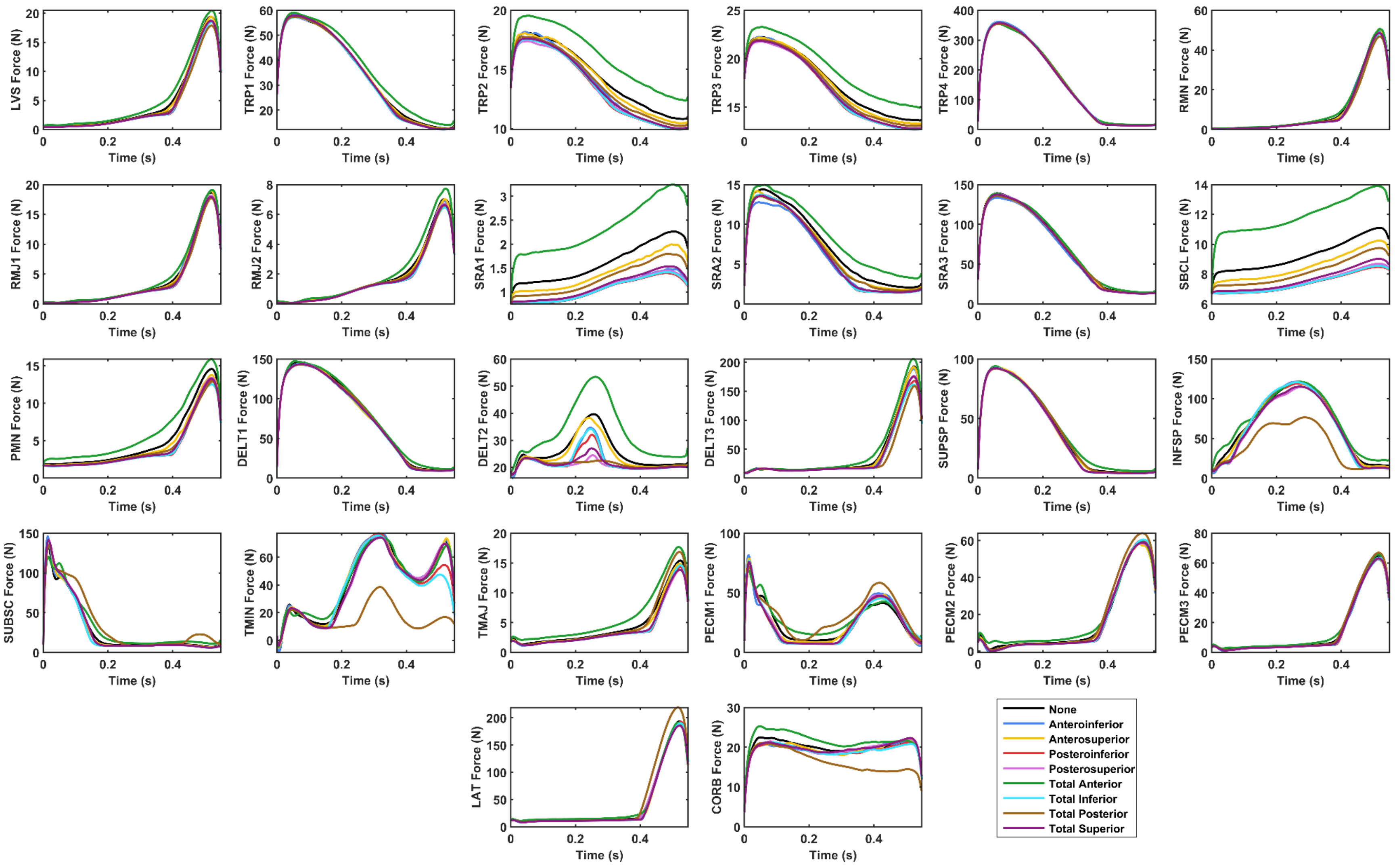


**Figure S8** **Muscle forces (in Newtons) of individual muscles for the forward reach task.**

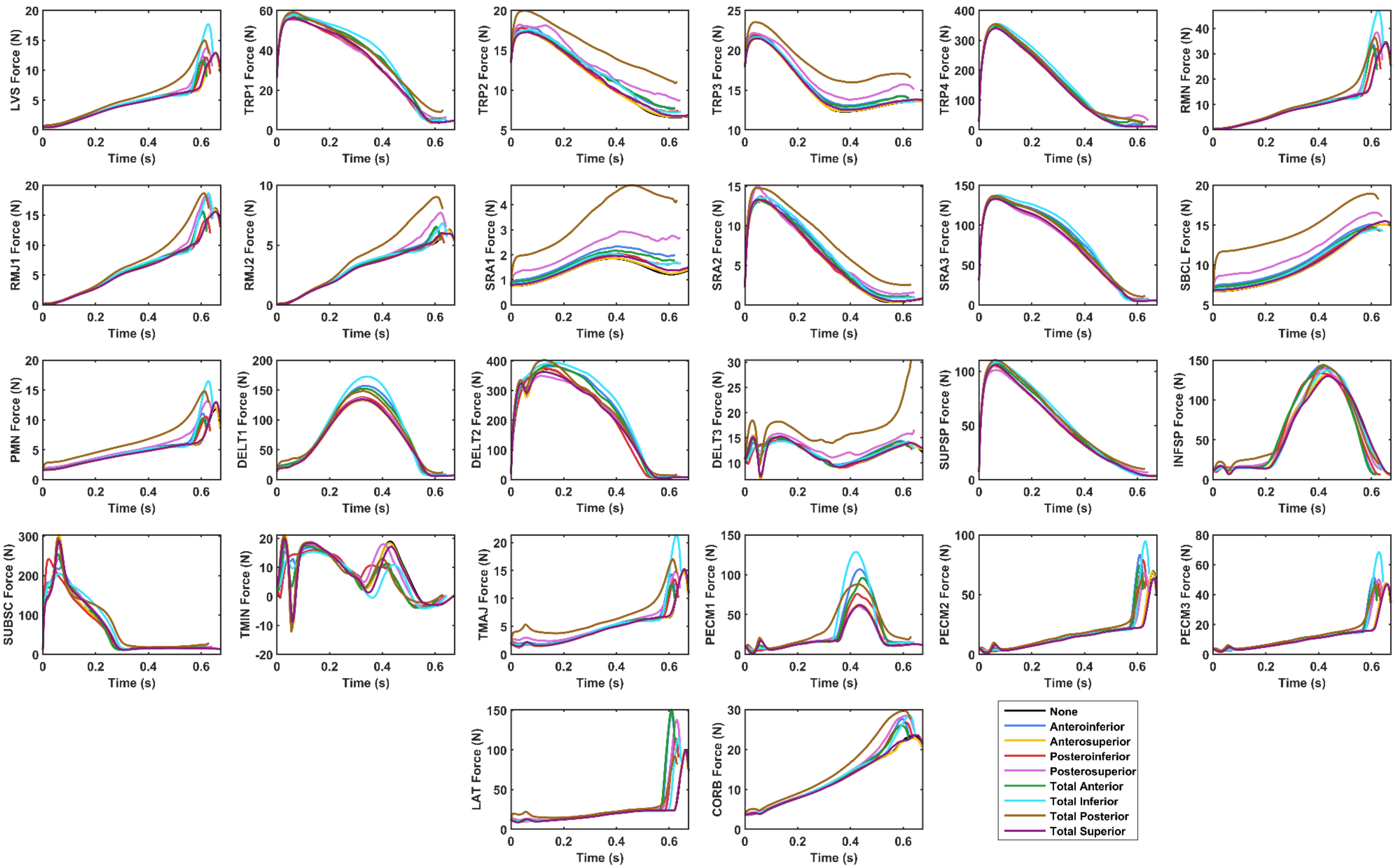


**Figure S9** **Muscle forces (in Newtons) of individual muscles for the head touch task.**

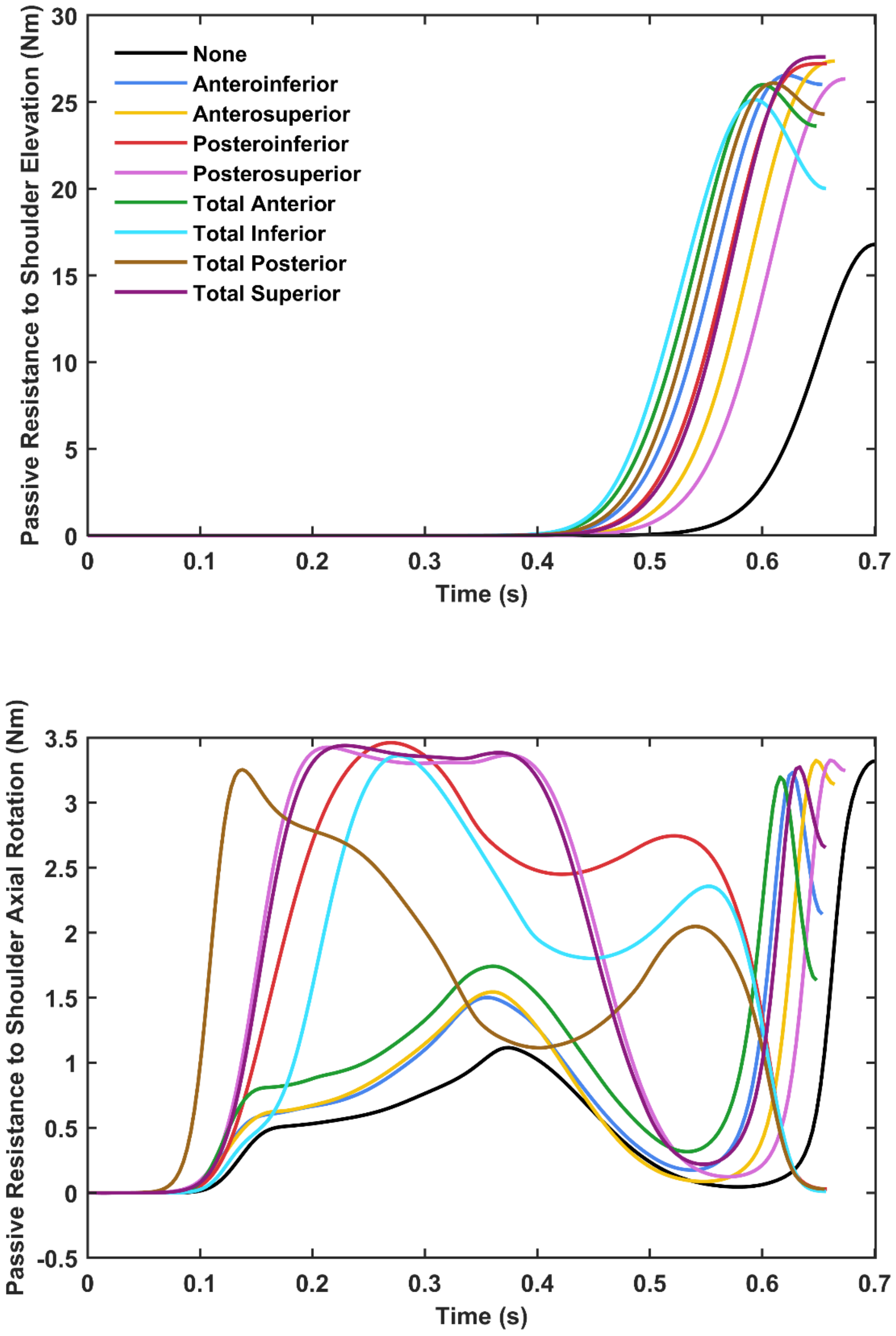


**Figure S10** **Torque generated by the simulated passive restraints during the upward reach task.**

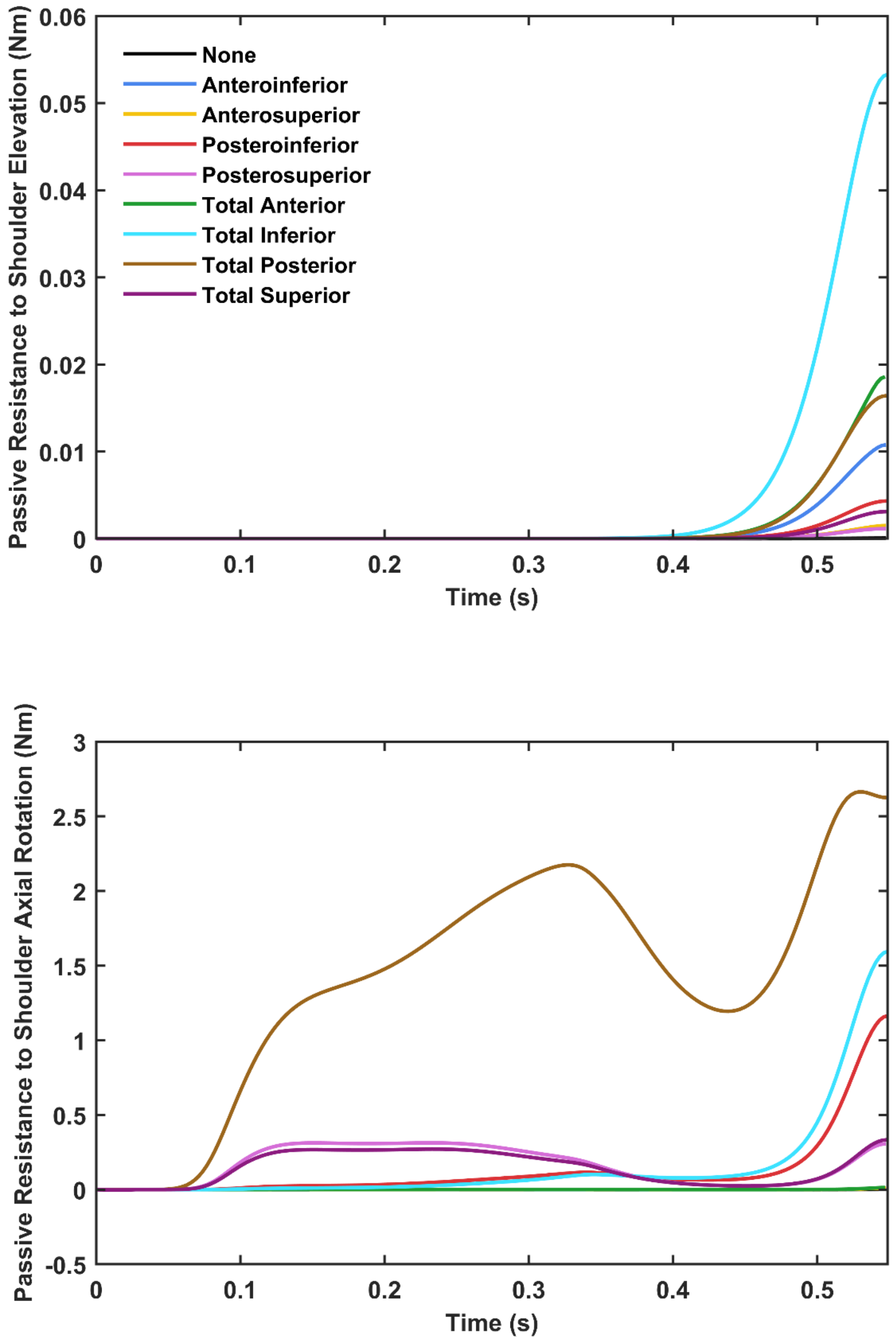


**Figure S10** **Torque generated by the simulated passive restraints during the forward reach task.**

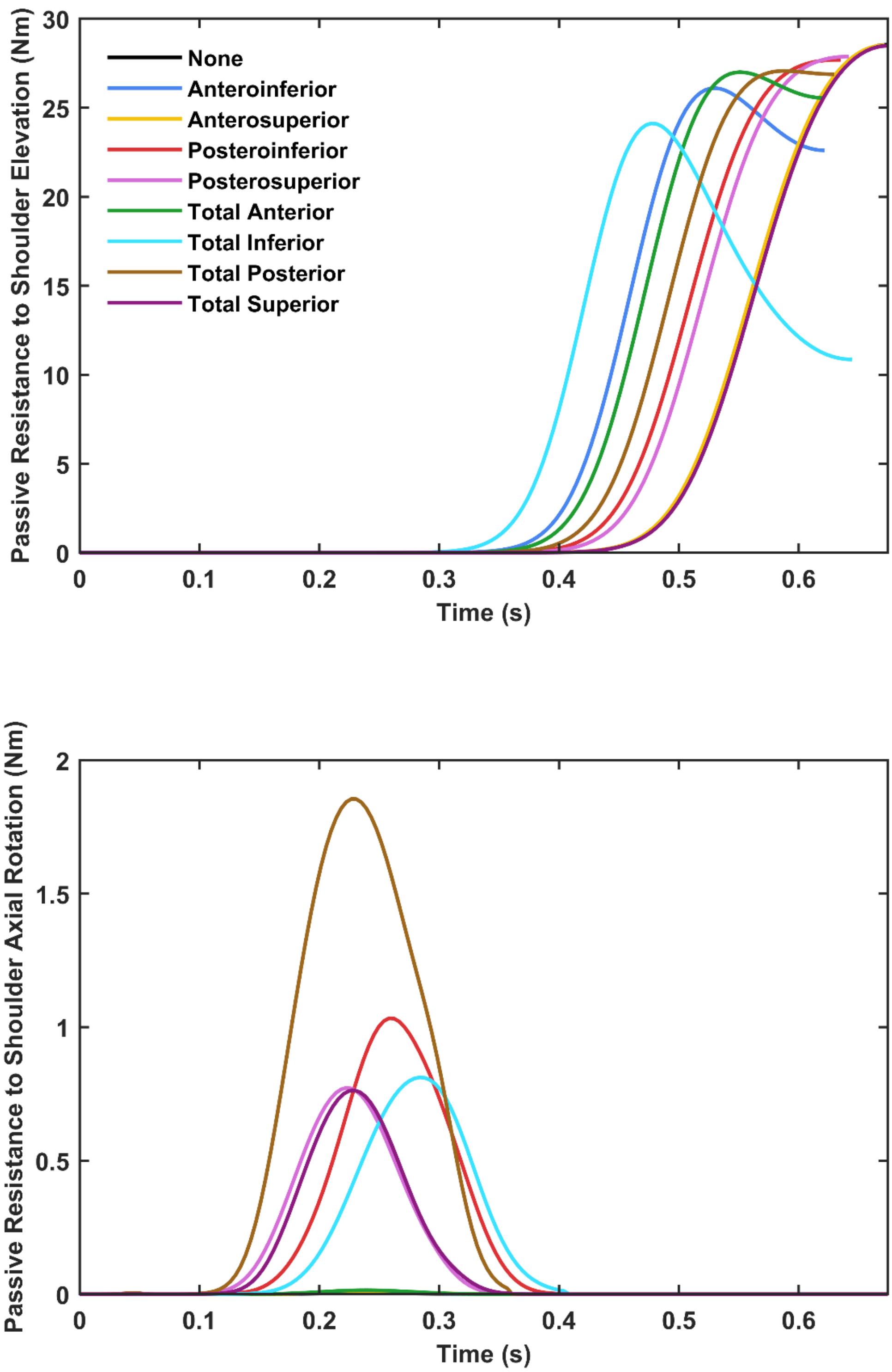


**Figure S10** **Torque generated by the simulated passive restraints during the head touch task.**
